# *Chop*/*Ddit3* depletion in β-cells alleviates ER stress and corrects hepatic steatosis

**DOI:** 10.1101/2020.01.02.893271

**Authors:** Jing Yong, Vishal S. Parekh, Jonamani Nayak, Zhouji Chen, Cynthia Lebeaupin, Jiangwei Zhang, Thazha P. Prakash, Sue Murray, Shuling Guo, Julio E. Ayala, Leslie S. Satin, Randal J. Kaufman

**Affiliations:** Degenerative Diseases Program, Sanford-Burnham-Prebys Medical Discovery Institute, 10901 N. Torrey Pines Rd., La Jolla, CA 92037; Department of Pharmacology, University of Michigan Medical School, 1000 Wall St., Ann Arbor, MI 48105; Department of Medicine, University of California San Diego, La Jolla, CA 92093; Department of Antisense Drug Discovery, Ionis Pharmaceuticals, Inc., 2855 Gazelle Court, Carlsbad, CA 92010; Cardiometabolic Phenotyping Core, Sanford-Burnham Medical Research Institute, 6400 Sanger Road, Orlando, FL 32827; Department of Molecular Physiology & Biophysics, Vanderbilt University School of Medicine, Nashville, Tennessee, USA

## Abstract

Type 2 diabetes (**T2D**) is a metabolic disorder characterized by hyperglycemia, hyperinsulinemia and insulin resistance (**IR**). During the early phase of T2D, insulin synthesis and secretion by pancreatic β cells is enhanced, which can lead to proinsulin (**ProIns**) misfolding that aggravates endoplasmic reticulum (**ER**) homeostasis in β cells. Moreover, increased insulin in the circulation may contribute to fatty liver disease. Medical interventions aimed at alleviating ER stress in β cells while maintaining optimal insulin secretion are therefore an attractive therapeutic strategy for T2D. Previously, we demonstrated that germline *Chop* gene deletion preserved β cells in high fat diet (HFD) fed mice and in leptin receptor-deficient *db/db* mice. In the current study, we further investigated whether targeting *Chop/Ddit3* specifically in murine β cells confers therapeutic benefits. First, we show that *Chop* deletion in β cells alleviates β cell ER stress and delays glucose-stimulated insulin secretion (**GSIS**) in HFD fed mice. Second, importantly, β cell-specific *Chop* deletion prevented liver steatosis and hepatomegaly in aged HFD fed mice without affecting basal glucose homeostasis. Third, we provide the first mechanistic evidence that ER remodeling secondary to *Chop* deletion modulates glucose-induced islet Ca^2+^ oscillations. Finally, using state-of-the-art GLP1-conjugated *Chop* AntiSense Oligonucleotides (GLP1-*Chop* ASO), we demonstrated that the *Chop* deletion induced GSIS change is a long term complex event in β cells. In summary, our results demonstrate that *Chop* depletion in β cells is a new therapeutic strategy to alleviate dysregulated insulin secretion and the consequently fatty liver disease in T2D.

---

Type 2 diabetes (**T2D**) is a metabolic disorder that poses a severe health challenge for modern society as it is estimated by the United States’ Centers for Disease Control and Prevention that thirty million Americans are affected by this condition (*1*). T2D is characterized by insulin resistance (**IR**), hyperglycemia and hyperinsulinemia (*2*). At the same time, current T2D therapeutics focus on achieving improved blood glucose homeostasis by improving both insulin secretion and reducing peripheral IR. Pharmacological interventions such as glitazones and glucagon-like peptide (**GLP1**) receptor agonists are limited in that despite achieving glucose control, there is insufficient clinical evidence to support a beneficial effect on human pancreatic β cells (*3*).

Pancreatic β cell pathogenesis coupled with peripheral IR has been the traditional explanation for T2D, although recent studies using mouse models and clinical findings made in the Pima Indians (*4*) support an alternative hypothesis that hyperinsulinemia can serve as a driving force for IR in mouse T2D models (*5, 6*). In the *Mehran and Johnson* paradigm (*5*), excessive amounts of insulin secreted by pancreatic β cells is the cause of peripheral IR and fatty liver development (*7*). In this model, reducing the insulin load would, therefore, alleviate IR. However, experimental observations made using murine *Insulin* gene KO models may not be relevant to human T2D, although a moderate reduction in insulin production may be beneficial (*7*). Nevertheless, it is technically challenging to accurately manipulate insulin mRNA levels (*8*), especially since it is the most abundant β cell mRNA, accounting for ~30% of the total transcriptome in mature β cells based on our own RNA-Seq data (*9*) and other’s findings (*10, 11*). Therefore, no current means exist to directly evaluate whether a reduction in insulin expression can achieve beneficial metabolic effects.

On the other hand, previous studies demonstrated that germline deletion of *Chop (also known as Ddit3/Gadd153)* prevents β cell failure in diabetes models (*12-14*). It is unknown, however, whether *Chop* deletion protects in β cells in a cell autonomous manner, especially considering the role of CHOP in reducing weight gain (*12*), regulating hepatic lipid metabolism and suppressing adipocyte development (*13*). To critically evaluate the metabolic consequence of deleting *Chop* in β cells, we therefore generated a conditional *Chop* deletion model by breeding a floxed *Ddit3* gene allele (*15*) with a *RIP-CreERT* transgene (*16*). In this model, β cell-specific *Chop* deletion is temporally controlled by tamoxifen (**TAM**) injections (referred to hereafter as “*Chop Δ/Δ: RIP-Cre*” or simply “*Chop βKO”* mice, see **Sppl. Table I** for breeding scheme).

To evaluate the primary effect of *Chop* deletion, we first assessed blood glucose and insulin levels in normal diet-fed male *Chop βKO* mice (with TAM administered to 21wk-old mice), compared to the isogenic male littermates that received diluent as controls (n=4 males/group). Four months after *Chop* deletion, *Chop βKO* mice had similar fasting blood glucose levels and displayed similar responses to a glucose challenge (*i.p.* at 1.5 mg glucose/g weight, Fig. 1A) compared to control littermates. Interestingly, fasting insulin levels were reduced, although not significantly, while glucose stimulated insulin secretion (**GSIS**) was dramatically blunted at 30min (Fig. 1B, p< 0.05 for 30min). In addition, we found a trend of reduced β-cell function, by applying the Homeostasis Model Assessment of β-cell function (**HOMA-β**) (**Fig. S1A**, p= 0.06), but not of HOMA-IR (**Fig. S1B**, p= 0.31) (*17*) suggesting unchanged IR in *Chop βKO* mice. Nonetheless, there was a significant linear correlation between fasting insulin and body weight (Fig. 1C, p< 0.01 by F-test). Furthermore, there was a trend towards decreased cumulative at 4 months (4mon) weight gain in *Chop βKO* mice (Fig. 1D, p= 0.11), which became significant when we included the non-*CreERT* littermates in comparison (**Fig. S1C**, p< 0.05). Importantly, cumulative weight gain as a function of time was not affected by *CreERT* gene expression as we previously reported (*9*) (and data not shown).

**Figure 1.**
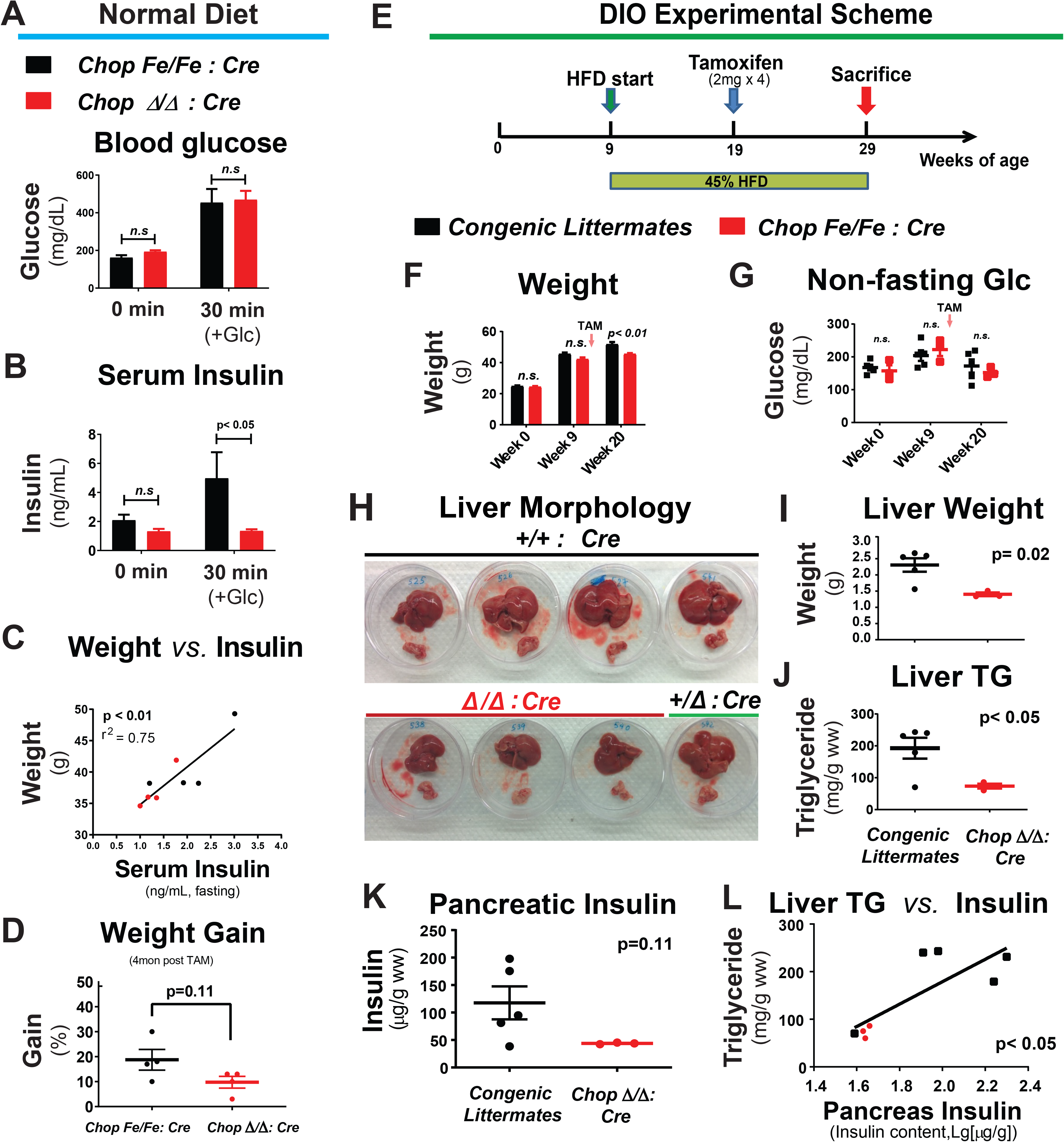
β-cell specific *Chop* deletion reduces pancreatic and circulating insulin levels and prevents liver triglyceride accumulation in male HFD fed mice. Under normal diet feeding, *Chop* floxed (n=4) and the isogenic *Chop* deleted (n=4) littermates were tested 4mon after TAM injections. After a glucose injection (1.5 mg/g body weight, *i.p.*), there was no difference in glucose excursion after 30min (**A**), while serum insulin levels were significantly reduced after 30min in *Chop*-deleted mice (**B**, p< 0.05, by Bonferroni’s test after RM 2-Way ANOVA). (**C**) The body weight of individual mice correlated linearly with fasting insulin levels (p< 0.01 by F-test, r^2^=0.75). (**D**) *Chop* deleted mice displayed a non-significant decrease in cumulative weight gain at 4mon after gene deletion (p=0.11 by 2-tailed *Student’s* t-test). In a separate experiment, (**E**) *Chop* floxed (n=3) and congenic control littermates (n= 5, with 4 WT animals +1 heterozygous animal) were fed high fat diet (HFD, 45% kcal from fat) for 20wks before tissue harvest. Mice received TAM injections at 10wks after HFD to delete *Chop* gene in β-cells. Experimental scheme is shown. (**F)** Reduced body weight gain of *Chop* deleted mice became significant at 20wks (p< 0.01, 10wks after *Chop* deletion by TAM). (**G)** No significant difference was detected for non-fasting blood glucose levels between the two groups. (**H)** Fresh liver (upper) and pancreas (lower) are shown for all mice, immediately after tissue dissection. (**I)** Liver weight was significantly less for *Chop*-deleted animals (*Chop Δ/Δ*: *Cre*, after CreERT-mediated *Chop* deletion) compared to WT and Het littermate control mice (p= 0.02). (**J)** Liver triglyceride levels were significantly lower in *Chop-*deleted mice, compared to WT and Het littermate control mice (p< 0.05). (**K)** Pancreatic insulin contents were reduced in *Chop* deleted mice compared to littermate control mice, albeit non-significantly (p= 0.11). (**L)** Liver triglyceride content correlated linearly with pancreatic insulin content on a semi-log plot (p< 0.05).

Given that male *Chop βKO* mice had relatively normal phenotypes, we next challenged them with a high-fat diet (**HFD**, 45% fat in kcal) for 20wks starting from an age of ~9 wks, with *Chop* deletion induced by TAM at 10wks post HFD (scheme shown in Fig. 1E). In addition, we selected littermates harboring wildtype (WT) *Chop* alleles as the control group as they can be TAM treated. Before *Chop* deletion, the two groups were metabolically indistinguishable with no significant difference in body weight or blood glucose levels, either before or 10wks after the HFD (Fig. 1 F&G). After *Chop* deletion, a moderate but significant decrease in body weight was observed in *Chop βKO* mice (Fig. 1F, p< 0.01 at *Wk20*), which had no effect on blood glucose (Fig. 1G). After 20wks HFD feeding, murine livers were dissected for visual inspection and for liver triglyceride (**TG**) analysis. HFD feeding caused hepatomegaly and liver discoloration associated with fatty deposits in control littermates (Fig. 1H, black line), as well as for a *Chop βHet* mouse in the litter (Fig. 1H, green line). In contrast, the three *Chop βKO* mice had normal-sized livers and appeared healthy (Fig. 1H, red line). The morphological impression was further confirmed quantitatively by showing significantly reduced liver weight and TG content in *Chop βKO* mice compared to littermates (Fig. 1 I and J, p< 0.05 for both). We surmised that chronically reduced *β* cell insulin secretion may prevent fatty liver development in the HFD-fed C57BL/6 mice, as previously proposed by others (*5, 6*). Supporting this hypothesis, pancreatic insulin content (standardized by tissue wet weight) was reduced 3-fold in the *Chop βKO* mice (Fig. 1K, p=0.11 due to variability in the control group) and was positively correlated with TG content (Fig. 1L, p< 0.05 by F-test). Echoing the recently reported human study (*7*), it will be of interest to investigate if the improvement of fatty liver we observed here reflects reduced *de novo* hepatic lipogenesis.

Intrigued by the blunted GSIS response and the protection from HFD-induced hepatic steatosis observed in the *Chop βKO* mice (compare Fig. 1B vs. Fig. 1H), we further evaluated the mice to test whether the reduced pancreatic insulin content negatively affected glucose metabolism. For this purpose, a follow-up 22 wk HFD experiment was performed using male mice, during which body weight, food intake and non-fasting blood glucose levels were all monitored. We found no significant differences in these variables at any time point when comparing knockouts to their *Chop βHet* littermates (Fig. 2 A to **C**, respectively), while glucose tolerance in all the HFD fed mice exhibited a significantly increased glucose excursion from baseline, as expected (*AUC-IPGTT*, Fig. 2D, p=0.0001 for “*HFD*” by 2-way ANOVA). In the same experiment, *Chop* βKO HFD mice were more sensitive to insulin only after *Chop* deletion, although the difference was not statistically significant (compare **Sppl. Fig. 2A** and **2B**, with TAM administered at 8wks post HFD). Similar to the insulin measurements made in the sera from mice fed normal diet (Fig. 1B), *Chop βKO* HFD fed mice had slightly decreased serum C-peptide, both before and after glucose stimulation (**Sppl. Fig. S2C**).

**Figure 2.**
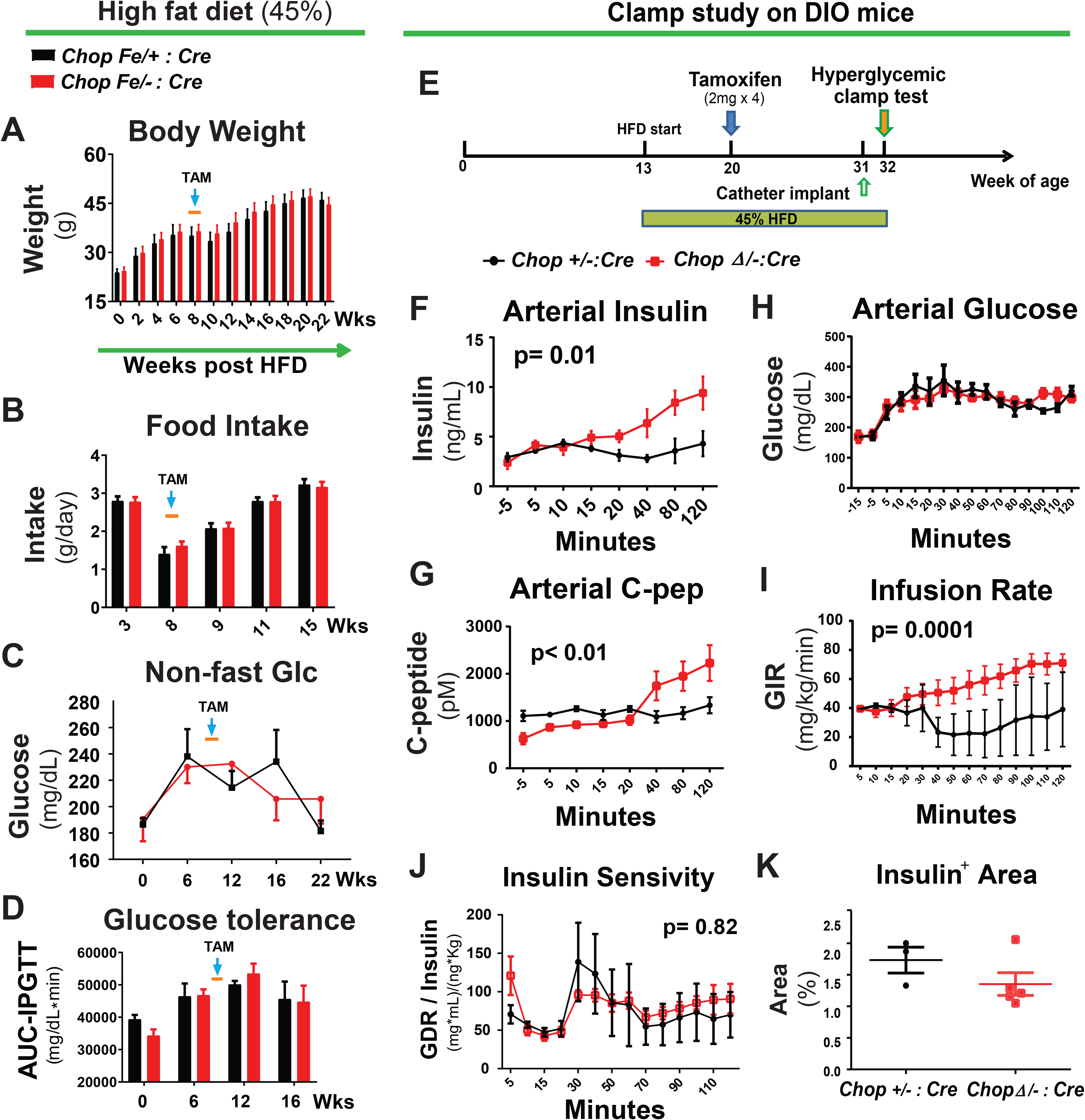
Beta-cell specific *Chop* deletion improves insulin secretion after sustained glucose stimulation, without altering glucose metabolism. Mice at 9wks of age were fed 45% HFD for 22wks and after 10wks TAM was injected. Floxed *Chop* heterozygous littermates served as controls (for which deletion of one allele was accomplished by TAM injection). There was no significant difference in body weight (**A**), daily food intake (**B**), non-fasting blood glucose levels (**C**) or glucose excursion after an IPGTT (**D**). In a separate experiment, similar comparison groups were used in a hyperglycemic clamp test: (**E**) shows Experimental Scheme. *Chop-*floxed mice with their congenic littermates were fed with HFD for 7wks before TAM injection. Mice were fed HFD for another 12wks before the hyperglycemic clamp assay (n=3 for *Chop +/-: Cre* and n=8 for *Chop Δ/-: Cre*, both male and female mice were tested). (**F and G)** Insulin and C-peptide ELISA assays demonstrated a significant increase in *Chop* deleted mice after 40min (p= 0.01). (**H)** The hyperglycemic clamp was maintained at 300 mg/dL by continuous glucose infusion into the jugular vein via an implanted catheter. (**I)** Glucose infusion rates (GIR) were significantly higher for β-cell specific *Chop* KO mice (p= 0.0001). **(J)** There was no difference in insulin sensitivity, defined as glucose disposition rate (GDR) divided by Insulin plasma concentration, between the two groups (p=0.82). After the hyperglycemic clamp, pancreata were dissected and processed for histological analysis. (**K)** Pancreatic insulin immuno-positive areas were similar in both groups. For panels **A to J**, RM-*2 Way ANOVA* was applied for statistics. Data are presented in dot plot for individual mice, with Bar and Whiskers representing Mean ± S.E.M.

As we observed no adverse effects of *Chop* deletion on mouse whole body metabolism, we performed a hyperglycemic clamp test on a different HFD cohort using mice of both sexes (Fig. 2E), again using age-matched *Chop βHet* littermates *(Chop +/-: RIP-Cre)* as controls. While no differences were observed in body weight nor fasting blood glucose (**Sppl. Table II**), it was intriguing to observe that both insulin and C-peptide secretion in *Chop βKO mice* appeared to be significantly slowed following hyperglycemia clamping, featuring a delayed, increased secretion after 40min (Fig. 2 F and **G**, p<= 0.01 by RM 2-way ANOVA labeled). Notably, the control *Chop βHet* mice showed basal hyperinsulinemia and reduced first phase responses to glucose demonstrating that the HFD model properly replicated the phenotype of humans during pre-diabetic phase and early T2D phase. *Chop βKO* mice required a significantly higher glucose infusion rate (**GIR**) to maintain their blood glucose target at ~300 mg/dL (=16.7 mM) (Fig. 2 H and **I**), suggesting increased glucose clearance was due to increased insulin secretion and not altered insulin sensitivity (p=0.82, Fig. 2J). Similarly, no difference was found in HOMA-β nor HOMA-IR (**Fig. S3 A and B**, p= 0.75 and 0.72, respectively). Furthermore, the differences found in the hyperglycemic clamp study were not due to altered *β* cell mass of the *Chop βKO* mice, as no statistical significance was found in the fractional insulin immuno-positive (*Insulin^+^*) areas (Fig. 2K, representative islet morphologies shown in **Sppl Fig. S4**, with total surveyed pancreas areas reported in **Sppl Fig. S5A**). Similarly, no differences were observed in the two groups with regards to *α* cell distribution in the islets (**Sppl. Fig. S4B** vs. **S4D**), *α* cell mass, or the relative ratio of α cells to *β* cells (**Sppl. Fig. S5B**-**D**). Finally, we ruled out the possibility of an indirect effect of *β-*cell *Chop* deletion on hepatic gluconeogenesis using a pyruvate tolerance test (**Sppl. Fig. S6**). In contrast, an *in vitro* GSIS assay using germline *Chop* KO islets confirmed a delayed, increased GSIS phenotype (**Sppl. Fig. S7**, p<0.01 for “*240 min*”), pinpointing an islet-autonomous change in GSIS accounting for our observations in whole animals.

To confirm that *Chop* deletion in *β-*cells reduces ER stress (*12*), we performed molecular analysis of UPR markers using qRT-PCR and subsequently using whole transcriptomic profiling by mRNA-Seq (*9, 18*) on the RNA extracted from the islets of male HFD fed mice. Molecular assays revealed greatly decreased *Insulin* transcripts (Fig. 3A, ~75% reduction in *Chop βKO* islets) associated with reduced UPR markers, for example, *Atf4*, *Bip* and spliced *Xbp1* (*sXbp1*) in *βKO* islets compared to *WT* littermates (Fig. 3B). Furthermore, supporting our findings by qRT-PCR, transcriptome profiling showed significant reduction in other UPR markers, represented by *Atf3, Bip/Hspa5, Sec23b* and *ERdj4/Dnajb9* on a “*Volcano plot*” (Fig. 3C). Interestingly, *Chop* deletion did not affect β cell identity, as the key β cell transcription factors, including *Pdx1, Nkx6.1, Mafa, Isl1* and *Ngn3*, were not altered in *Chop*-deleted islets (Fig. 3D), consistent with our histological findings of unchanged β-cell mass and islet morphology. In addition, *Chop* deletion did not alter the mRNAs encoding *Cpe, Glut2* or *PC1/2* (**Sppl. Fig. S8**), further supporting the hypothesis that *Chop* deletion did not impair β cell differentiation. In contrast, *Wfs1*, *Gadd34/Ppp1r15a*, and *Bip*/*Hspa5* (encoded by two alternatively spliced isoforms, namely *NM_022310* and *NM_001163434*), representing the UPR target genes for the IRE1α, PERK and ATF6α branches of the UPR, respectively, were all found to be significantly reduced (Fig. 3E). Reductions in UPR transcript abundance was supported further by our finding that a subset of mRNAs that encode important ER proteins were altered (see heat map in Fig. 3F), suggesting that CHOP has a long-term effect on “*ER remodeling*”. Since the UPR is an adaptive response to ER stress, we tested biochemically if *Chop* deleted β cells exhibit reduced ER stress, by probing the interaction between ProIns and BiP, as BiP binding to misfolded proinsulin is the gold-standard biochemical indicator for ER due to ProIns misfolding in the β cell (*19-21*). By the BiP co-IP assay, we confirmed that less BiP protein was associated with ProIns in the *Chop* β*KO* islets (Fig. 3G, and data not shown), confirming reduced ER stress in *Chop* β*KO* islets.

**Figure 3.**
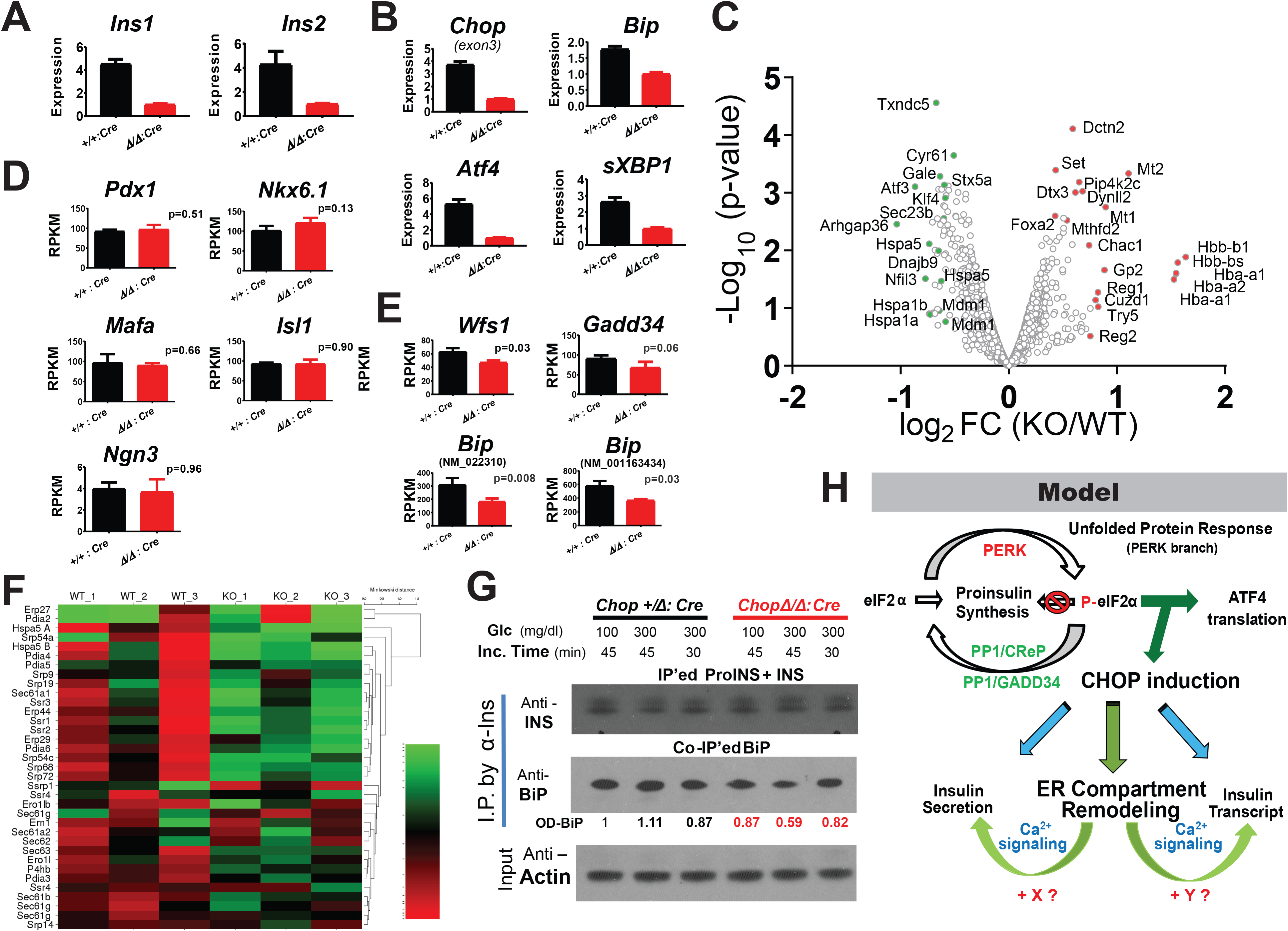
*Chop* deletion reduces *Insulin* transcripts and reduces ER stress in islets of male HFD fed mice. The *Chop* floxed (*Fe/Fe: Cre*) and their wildtype *(+/+:Cre*) congenic littermates (n=5/group, all male) were fed HFD (45% cal from fat) for 45wks before TAM injections and were continued on HFD for another 8wks before islet isolation. Transcripts encoding *Insulin* genes (*Ins1* and *Ins2*) and UPR genes were reduced in *Chop KO* islets. Representative qRT-PCR results from one pair shown (**A**, **B)**. The same qRT-PCR experiment was repeated five times using different pairs of mice, e.g. results shown in Fig. S9. Subsequently, islet RNA was extracted from the same donors as in **A** and **B** were further analyzed by RNA-Seq (n=3/ group). Transcriptome profiles were performed using poly-A enriched mRNA by Next-Generation Sequencing. (**C)** Volcano plot of mRNA differential expression was generated for both groups. Genes with high fold-change and/or highest p-values were labeled as green/red dots to reflect down-/up-regulation as a result of *Chop* deletion. (**D)** RPKM values for important β-cell lineage genes *Pdx1, Nkx6.1, Mafa, Ngn3 and Isl1*, were unchanged and plotted with p-value indicated for comparison between KO and WT mice. (**E)** RPKM values for selected UPR genes were plotted with p-values indicated. (**F)** Heat-map was generated for a selected group of genes important for ER protein synthesis/translocation, with percentage change represented by the heat map. (**G)** Co-IP verified decreased ER stress in β-cells. BiP-bound ProINS was quantified by anti-Ins co-IP in β-cell specific *Chop* KO islets, 6wks after TAM injection. Islets from male donors (*Chop Δ/+: Cre vs. Chop Δ/Δ: Cre*, n=3 /group) were pooled for overnight culture an then glucose challenged. Two concentrations of glucose were tested as indicated. INS and ProINS proteins were IP’ed by a monoclonal mouse anti-INS antibody (Invitron, Clone 3B1) and BiP was detected by a monoclonal rabbit anti-BiP antibody (kind gift from L. Hendershot). Co-IP’ed BiP was further quantified by optical density (OD). (**H)** Working Model: *Chop* induction calibrates ER stress from ProIns synthesis. In turn, CHOP mediates ER remodeling to influence insulin transcription and granule release, mainly through ER Ca^2+^ signaling (indicated by green arrowheads) and likely through other metabolic cues (unidentified, represented by “X” and “Y”).

These results led us to hypothesize that under physiological conditions, CHOP serves as a transcriptional hub in β cells that can alter ER function by reducing the expression of genes encoding ER-structural, -functional and ER-to-Golgi trafficking proteins in order to accommodate ProIns synthesis (Fig. 3H, Model). Conversely, *Chop* deletion in β cells decreased ProIns synthesis accompanied by more efficient folding as indicated by less BiP interaction.

A missing mechanistic link, however, is that CHOP does not directly bind to the promoter regions of all of these genes affected (Fig. 3F), as we demonstrated previously using ChIP-Seq (*18*). We thus wondered how deleting *Chop* could result in changes in ER function without altering the promoter activity of ER-related genes. The ER is the major intracellular Ca^2+^ storage organelle, and both ER and cytosolic Ca^2+^ and ATP-to-ADP levels can be profoundly affected by physiological ER stress (*22, 23*). Ca^2+^ has a crucial physiological role in pancreatic islets as glucose-dependent cytosolic Ca^2+^ oscillations driven by membrane electrical activity trigger insulin exocytosis (*24-26*) and *Insulin* gene transcription is positively regulated by cytosolic Ca^2+^ levels (*27, 28*). We therefore tested whether the *Chop* deletion-dependent changes in islets were associated with altered ER Ca^2+^ signaling, representing a logical outcome from reduced ER stress (Fig. 3).

For this purpose, we tested the effect of a membrane-permeable intracellular Ca^2+^-chelator, BAPTA-AM (abbreviated as “BAPTA” hereafter). When islets isolated from C57BL/6 mice were exposed overnight to BAPTA (10 µM), we observed reductions in both *Ins1* and *Ins2* mRNA (Fig. 4A, “*Insulin*” panel, *Ctrl* versus *BAPTA*). At the same time, *Chop* and *sXbp1* levels were also reduced by BAPTA treatment, suggesting that lowering β-cell Ca^2+^ reduced UPR induction in these *WT* islets, possibly by decreasing the ProIns translation burden placed on the ER (Fig. 4A, “*Chop*” and “*Xbp1*” panel, *Ctrl* columns versus *BAPTA-AM*columns). Importantly, tunicamycin (**Tm**, 0.5 µg/mL) in combination with BAPTA treatment rescued the decrease in both *Ins1* and *Ins2* mRNA levels, showing that the effect of Ca^2+^ chelation on β-cell ER stress could be overcome by artificial “ER stress” induction. Concomitantly, *Chop* and *sXbp1* mRNAs, as well as *BiP* mRNA were positively induced by Tm treatment (Fig. 4A, *BAPTA* columns versus *BAPTA+Tm* columns).

**Figure 4.**
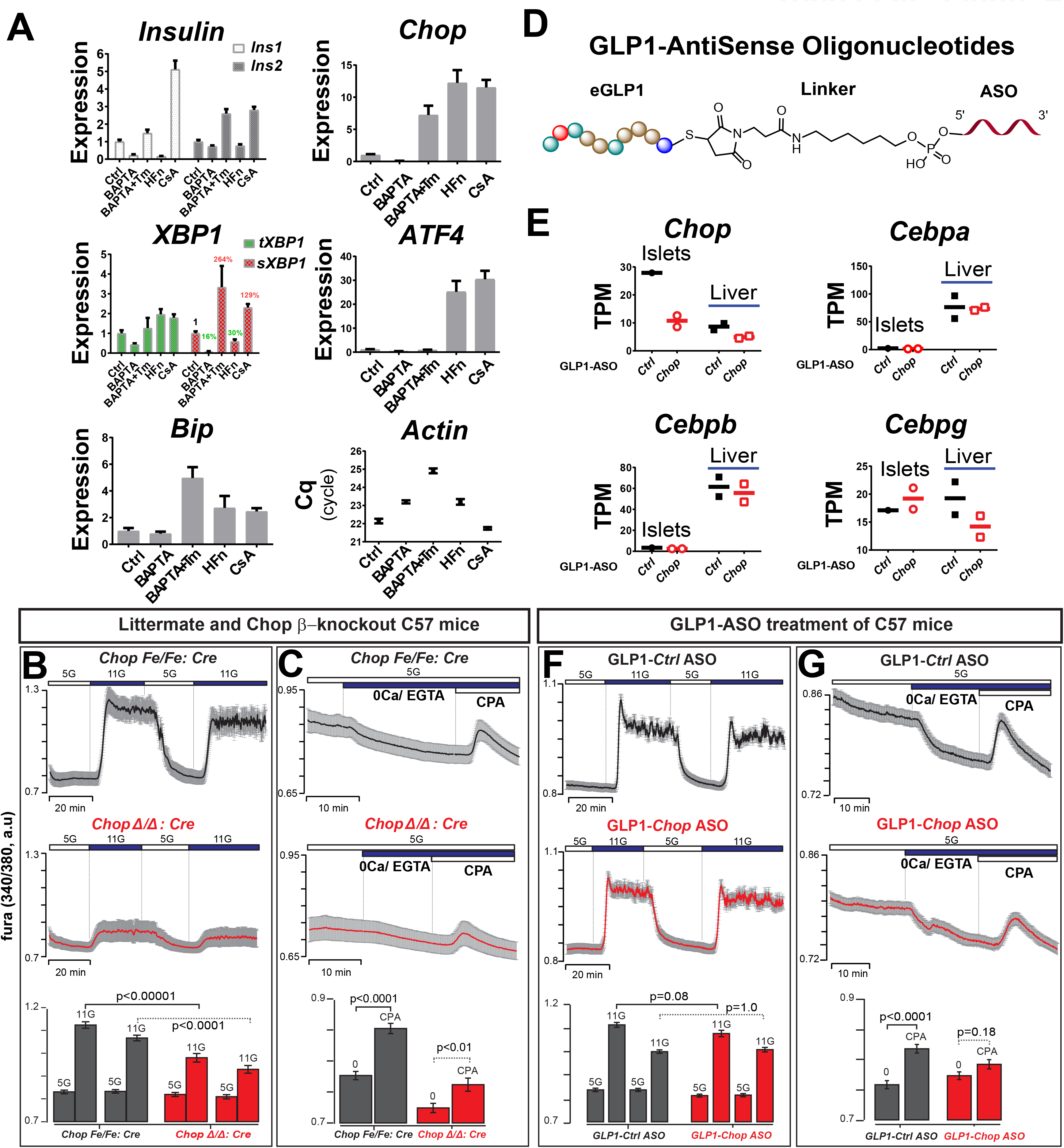
ER stress induces insulin mRNAs via increasing the releasable ER Ca^2+^ pool and *Chop* depletion reduces the ER Ca^2+^ pool. **(A)** Islets isolated from male C57Bl6/J (n=5) were pooled and divided into five groups (after overnight culture, with ~120 islets/group). The islets were subsequently challenged with either DMSO (labeled as “*Ctrl*”) or one of the following compounds: 1) 10µM *BAPTA-AM*; 2) 10µM *BAPTA-AM* plus 500ng/mL *Tm*; 3) 50nM halofunginone (*HFn*); and 4) 200nM *CsA*, for 18hrs before RNA extraction for qRT-PCR. The relative transcript abundance is summarized by bar graphs, using *β-Actin* expression (expressed as “Mean Cq”) as a reference. Both cytosolic Ca^2+^ (**B**) and ER Ca^2+^ pools (**C**) were reduced in β-cell specific *Chop* KO islets. Representative traces show intracellular Ca^2+^ signal (mean ± 95% CI) (n = 30-50 islets/group). Experiments were repeated three times using three pairs of control and β-cell *Chop*-deleted islets. Experimental conditions included physiological extracellular Ca^2+^ (2.56mM), followed by extracellular Ca^2+^ removal, and exposure to the reversible SERCA pump inhibitor CPA (50μM) with glucose at 5mM (“5G”) or 11mM (“11G”). Two-way ANOVA with post-hoc Tukey multiple comparison test were used for statistical analysis. (**D)** Illustration shows GLP1-ASO chemical structure. (**E)** GLP1-*Chop*-ASO reduces *Chop* mRNA (expressed as TPM) specifically in murine islets compared to liver tissue. RNA extracted from the livers of the same mice served as internal controls for *Chop* expression, for RNA-Seq. Expression of additional *C/EBP* transcription factor family members are shown as controls. GLP1-*Chop-*ASO had no obvious effects on glucose-induced cytosolic Ca^2+^ level (**F**), and reduced ER Ca^2+^ pools (**G**) in primary islets *in vivo*. Representative intracellular Ca^2+^ traces (mean ± SEM) are shown (n = 30-50 islets/group). For panels **F** and **G**, male and female *Chop* floxed mice (with no *RIP-CreER* gene) were injected I.P. with control or GLP1-*Chop*-ASOs 8 days before islet isolation. Two-way ANOVA with post-hoc Tukey multiple comparison test was used for statistical analysis.

The positive regulation of *Ins1/Ins2* transcription by ER stress, previously reported (*9*), was further confirmed in a separate batch of *WT* and *Chop βKO* islets treated with 5 µg/mL Tm (**Sppl. Fig. S9**). In contrast, induction of *Chop* mRNA by activating GCN2 using halofunginone (**HFn**, 50nM), a potent tRNA synthetase inhibitor (*29, 30*) to induce eIF2α phosphorylation, actually reduced *Ins1* and *Ins2* transcripts, as well as sXBP1 in spite of pronounced *Atf4* and *Chop* induction (Fig. 4A, *Ctrl.* Versus *HFn*). Under HFn treatment, *Chop* induction is partially attributable to the amino acid response element (**AARE**) sequence in the *Chop* promoter (*31*). Lastly, addition of cyclosporine A (**CsA**, 200nM) to islet media induced both *Ins1* and *Ins2* transcripts, suggesting the ER Ca^2+^ effect on *Ins1/Ins2* expression was unlikely due to a direct consequence of activation of the Calcineurin/NFAT pathway (*32*) by Ca^2+^.

Insulin granule secretion from mature β cells is dependent on cytosolic Ca^2+^ oscillations and *Chop* β*KO* islets exhibited reduced cytosolic Ca^2+^ concentration in response to 11 mM glucose (Fig. 4B, p< 0.0001). Interestingly, ER Ca^2+^ content was also reduced in these islets (Fig. 4C). Furthermore, GLP1-conjugated antisense oligonucleotides (GLP1-ASO) demonstrated efficient gene knockdown in rodent islets (*33*), providing an attractive, independent strategy to mediate *Chop* knockdown to test our hypothesis. We therefore treated mice having floxed *Chop* alleles (*15*) (on the *C57BL/6* background without *CreERT*) with GLP1-*Chop*-ASO *in vivo* (Fig. 4D, GLP1-*Chop*-ASO), by subcutaneous injections in young mice of both sexes (0.5 nMole/g body weight, twice over 5 days, in the back of neck), with GLP1-conjugated control ASO (sequence not homologous to any known gene in the murine genome). As expected, GLP1-*Chop*-ASO administration was well tolerated by the mice and specifically reduced islet *Chop* transcript by >60% (Fig. 4E), with minimal effects on other C/EBP family members (i.e. *Cebpa*, *Cebpb* and *Cebpg*) in either islets or liver tissues, demonstrating β cell selectivity as reported (*33*). Further supporting our hypothesis, treatment with GLP1-*Chop*-ASO reduced the ER Ca^2+^ pool without secondarily affecting the cytosolic Ca^2+^ levels stimulated by 11 mM glucose (compare Fig. 4F and 4G), suggesting the change in ER Ca^2+^ was a primary event that immediately followed *Chop* knockdown. To our knowledge, this is the first proof-of-principle that GLP1-ASO strategy can be exploited to alter islet physiology and Ca^2+^ dynamics. These results support our working hypothesis featuring an ER-centric role of CHOP in pancreatic β cells (Fig. 3H, Model).

In summary, we propose that CHOP is activated by UPR signaling through the PERK branch, as a response to increased ProIns synthesis and misfolding (*34*). In turn, CHOP serves as a transcriptional hub to maintain ER proteostasis. This process also controls *Insulin* transcription, partially via Ca^2+^ signaling, from the ER to the nucleus, although this is unlikely directly mediated by CHOP (Fig. 3H, indicated by the blue arrow on right). Furthermore, ER remodeling in turn has a profound effect on ER Ca^2+^ that subsequently contributes to controlling the increase in cytosolic Ca^2+^ that occurs in response to elevated glucose (Fig. 3H, as indicated by the blue arrow on left). At the same time, comparison of genetic *Chop* deletion model versus the GLP1-ASO mediated *Chop* knockdown model demonstrated that *Chop* deletion induced GSIS change is a long term complex event in β cells, with the ER Ca^2+^ pool change preceding insulin mRNA reduction and GSIS decrease.

As the pancreatic islet is a nutrient sensing organ, and β-cells secrete insulin to increase anabolic metabolism upon nutrient availability, we speculate that when there is a surfeit of nutrition, increased insulin secreted into the circulation exacerbates IR and promotes fatty liver in humans (*7*), although there was a dissociation between HOMA-IR (*17*) and hyperinsulinemia in *Chop βKO* mice (**Fig. S1B and S3B**). We were initially intrigued to find that a 75% reduction in *Insulin* mRNA by *Chop* deletion did not produce a strong metabolic phenotype in mice, although this was not unprecedented (*5, 6, 11*). Insulin mRNA may be in excess in β-cells and crucial for glucose sensing. This may be better explained by a model (Fig. 3H) proposing that ER “stress” due to increased proinsulin synthesis is coupled to insulin secretion, mediated through an “ER Ca^2+^” response in β-cells. In the UPR signaling cascade, however, *Perk* deletion (*35*), eIF2α phosphorylation site mutation (*36, 37*), *Ire1α* deletion (*9*), *sXbp1* deletion (*38*) and *Atf6α* deletion (*39*) all caused deleterious outcomes in β-cells and were thus unsuitable therapeutic targets. Uniquely, *Chop* deletion may be the only example that can safely reduce “ER stress” in β-cells (*12*), by exemplifying a “thrifty gene” providing an evolutionary advantage during famine (*40*). This hypothesis is more attractive given our finding that β-cell specific *Chop* deletion prevented HFD-induced hepatic steatosis (Fig. 1H), echoing recent human findings (*7*). Inspired by the unique phenotype of *Chop*-deleted β-cells, we discovered that a GLP1-conjugated *Chop* ASO could partially recapitulate “ER remodeling” characterized by a reduced ER Ca^2+^ pool, thereby providing a promising new therapeutic strategy for further pharmacological characterization and refinement to combat human T2D and fatty liver disease.

## Acknowledgements

The generation of a *Ddit3* gene floxed mouse model was a collaborative effort with Dr. Ira Tabas at Columbia University. Drs. Jian-Liang Li, Feng Qi and Jun Yin at SBP Applied Bioinformatics core provided guidance and helpful discussion on the transcriptomic data analysis. Ms. Guillermina Garcia at the SBP Histology Core facility provided technical assistance and helpful discussion on histology quantification methodology using the *Aperio Scanscope FL^®^* instrument. Tissue analysis was facilitated by the SBP Histology core at Lake Nona, with technical assistance from Mr. John Shelley.

R.J.K. is supported by NIH grants R01DK113171, R24DK110973, R37DK042394, R01CA198103 and the SBP NCI Cancer Center Grant P30 CA030199. R.J.K. is a member of the UCSD DRC (P30 DK063491) and an Adjunct Professor in the Department of Pharmacology, UCSD. L.S.S. is supported by NIH grant R01DK46409. C.L. is a member of the UCSD DRC and is supported by the NIH training grant T32DK007494. V.S.P. acknowledges support from an Upjohn Foundation postdoctoral fellowship.

