## Supplemental Document for "*Chop*/*Ddit3* depletion in β-cells alleviates ER stress and corrects hepatic steatosis"

Jing Yong et al.

### Supplemental Figures

#### Supplemental Figure 1

**Beta-cell specific *Chop* deletion in mice reduced β-cell function and cumulative body weight gain of male mice on normal diet, without affecting whole body IR.**

For the same groups of mice as shown in **Fig. 1** (**A** to **D**), (**A**) Beta cell function in Chop βKO mice was reduced with a boundary p value (p= 0.06 by unpaired two-tailed t-test), expressed in HOMA-β. (**B**) IR in Chop βKO mice was unchanged (p=0.31 by unpaired two-tailed t-test). (**C**) *Cho*p deletion in β-cells reduced cumulative body weight gain when littermates without the RIP-CreERT transgene were included for comparison (p< 0.05 by *Student*'s t-test).

#### Supplemental Figure 2

**β-cell specific *Chop* deletion in mice improved insulin sensitivity with reduced serum C-peptide,**

For the same groups of mice as shown in **Fig. 2** (**A** to **D**), insulin sensitivity and serum C-peptide levels were measured by ELISA at the indicated times. (**A**). At 5wks after HFD (**before** TAM injections), the two groups had identical insulin sensitivity in response to exogenously administered human insulin, at 1mU/g of body weight. (**B**). At 18wks after HFD (**after** TAM injections), *Chop* deleted mice displayed slightly improved insulin sensitivity (p=0.15 by 2-Way ANOVA R.M.). Data are presented as % of fasting glucose levels for individual animals, in the Mean ± S.E.M format. (**C**) In addition, at 20wks after HFD, *Chop* deletion reduced circulating C-peptide levels under fasting and glucose-challenged states (30min after a bolus injection of glucose at 1.5 mg/g). Data are presented as dot plots plus Mean ± S.E.M. Blood samples were drawn via retro-orbital bleeding.

#### Supplemental Figure 3

**Beta-cell specific *Chop* deletion in HFD fed mice did not change β-cell function and whole body IR**

For the same groups of mice as shown in **Fig. 2** (**E** to **K**), (**A**) Beta cell function in HFD fed Chop βKO mice mice was unchanged (p=0.75 by unpaired two-tailed t-test), expressed in HOMA-β. (**B**) Insulin resistance in in HFD fed Chop βKO mice was unchanged (p=0.72 by unpaired two-tailed t-test).

#### Supplemental Figure 4

**Islet morphology and distribution was not affected by β-cell specific Chop deletion.**

After the hyperglycemic clamp assay shown in **Fig. 2** (**E** to **K**), the same batch of mice were sacrificed with pancreata dissected and fixed for histological analysis. Representative images are shown for insulin positive areas in pancreatic sections, in control (**A** and **B**) and β-cell specific *Chop* deleted (**C** and **D**) mice. Sections were counterstained with H&E (**A** and **C**), or with DAPI for immunofluorescence (**B** and **D**), respectively. Microscopic images were captured by Aperio software.

#### Supplemental Figure 5

***Chop* deletion did not alter the α-cell to β-cell ratio nor the absolute Insulin/Glucagon positive areas.**

(**A**) For the same batch of mice used in **Fig. 2** (**E** to **K**), there were no differences in total pancreatic areas surveyed for the two groups. (**B**) Glucagon$^{+}$ *versus* Insulin$^{+}$ ratio by immunofluorescence staining in Chop β-KO pancreases was not significantly different from their heterozygous control littermates. There was no significant difference in the absolute areas for Glucagon+ areas (**C**) and for Insulin$^{+}$ areas (**D**).

#### Supplemental Figure 6

**Hepatic gluconeogenesis was not altered by *Chop* deletion in β-cells.**

Hepatic gluconeogenesis was tested using another independent set of male mice fed HFD for 22wks. A single bolus of 2 mg/g pyruvate was i.p. injected to increase glucose production via liver gluconeogenesis. No difference was detected in *Chop* β-KO mice (n= 3) versus their male littermates containing at least one allele of active *Chop* gene (n= 6). Blood glucose levels were measured at the indicated time points. Data are presented as Mean ± S.E.M.

#### Supplemental Figure 7

**Increased insulin secretion was delayed in germline *Chop* null islets.**

Islets freshly isolated from germline *Chop* heterozygous (**A**, *Chop Het*) mice and *Chop* null (**B**) littermates were morphologically indistinguishable. Scale bar = 400µm. (**C**) Islets were subjected to glucose stimulation (300mg/dL, ~16.7mM) for 20min and 240min, respectively, no difference was detected in insulin secretion at 20min after glucose stimulation. After prolonged glucose stimulation (300mg/dL x 240min), *Chop* null islets secreted more insulin ($p<0.01$, panel **C**). For panel C, aliquots of the incubating medium were taken for insulin ELISA assay at the end of the indicated incubation time.

#### Supplemental Figure 8

**β-cell specific *Chop* deletion did not alter transcripts encoding Glut2 and insulin processing enzymes measured by whole islet RNA-Seq.**

RPKM values (Mean ± SEM, n=3/group) for mRNAs encoding *Glut2* and insulin processing enzymes *Cpe*, *Pcsk1* and *Pcsk2* were plotted with p-value indicated for the comparison between β-WT and β-KO mice.

#### Supplemental Figure 9

**Tunicamycin treatment overnight increased *Ins2* transcript in primary islets.**

For the indicated β-WT and β-KO genotypes, islets from two animals were isolated, cultured overnight and pooled, before they were treated for 22hrs with Tm (5µg/mL) or DMSO as diluent control. Total islets RNA was extracted via *Trizol* method followed by Qiagen column purification. Transcript levels of Ins1 (**A**), Ins2 (**B**) and Gcg (**C**) were quantified by qRT-PCR, with the "*Expression*" levels standardized using *β-Actin* transcript abundance (also known as ΔΔCq). Specifically, the Tm-reduced *Ins1* transcript in β-WT islet samples may be an effect of excessive ER stress in this group.

**Yong et al., Sppl. Figure S1**

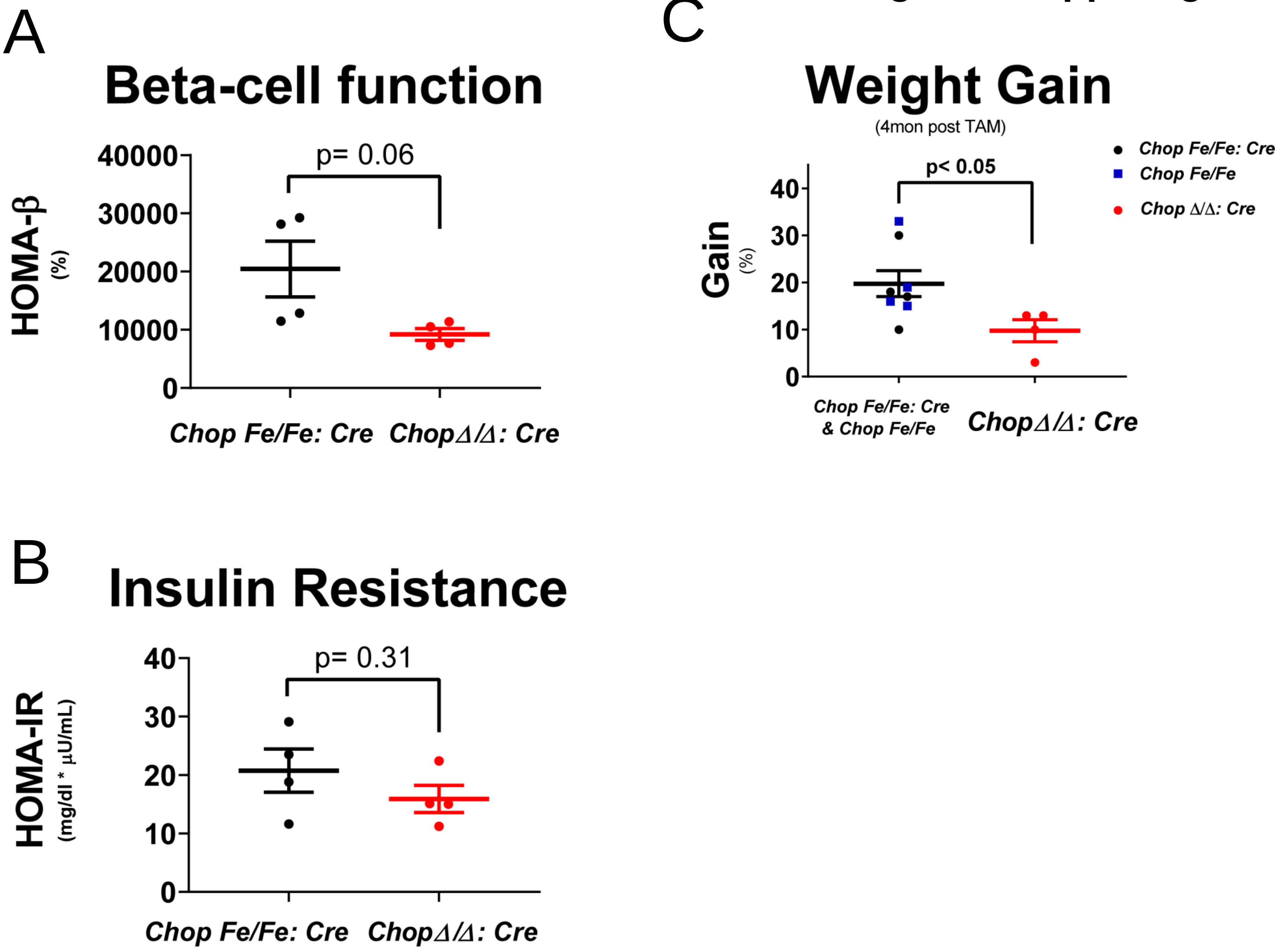

Yong et al., Sppl. Figure S2

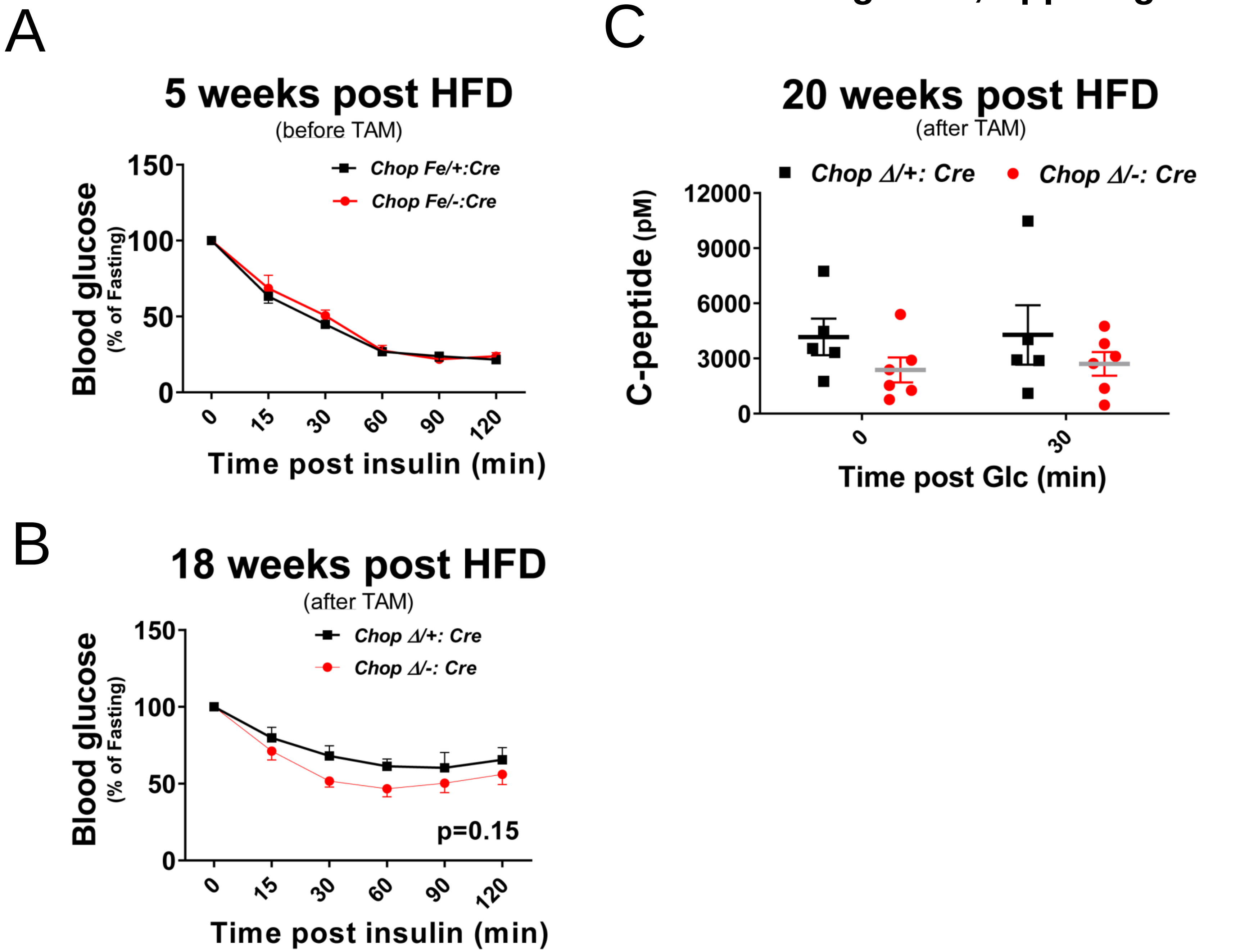

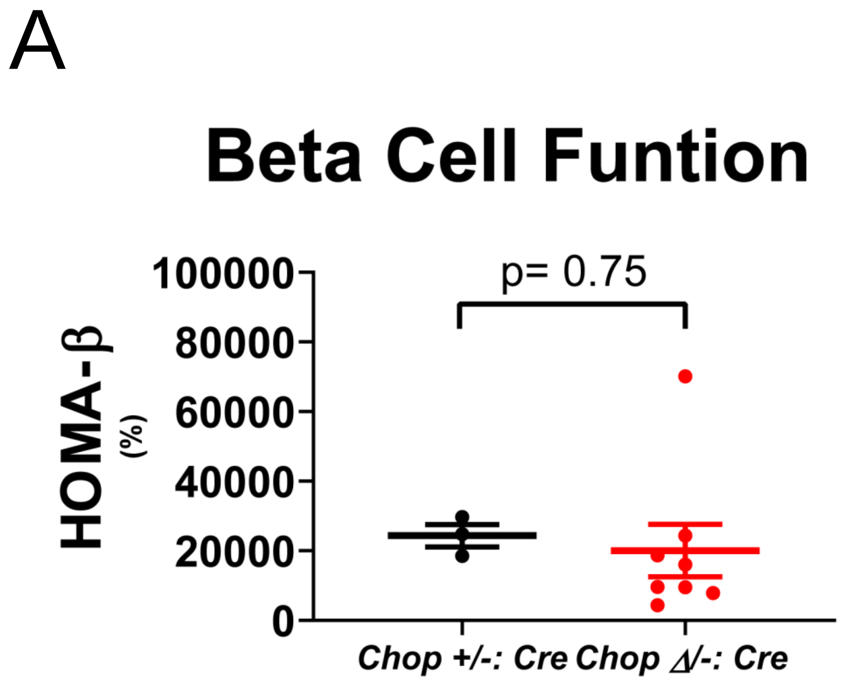
A
Beta Cell Funtion
p= 0.75
HOMA-β
(%)
100000
80000
60000
40000
20000
0
Chop +/-: Cre
Chop Δ/-: Cre

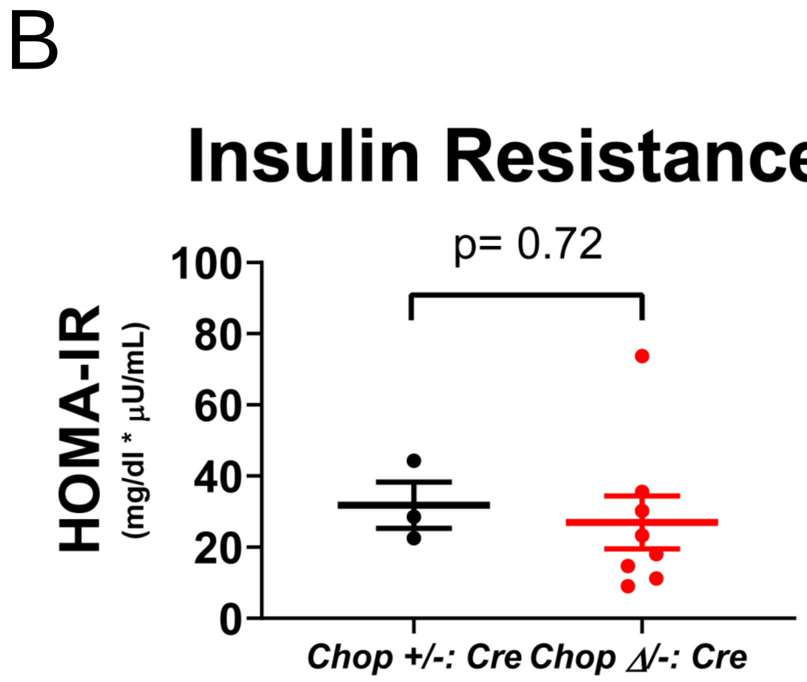
B
Insulin Resistance
p= 0.72
HOMA-IR
(mg/dl * μU/mL)
100
80
60
40
20
0
Chop +/-: Cre
Chop Δ/-: Cre

Yong et al., Sppl. Figure S4

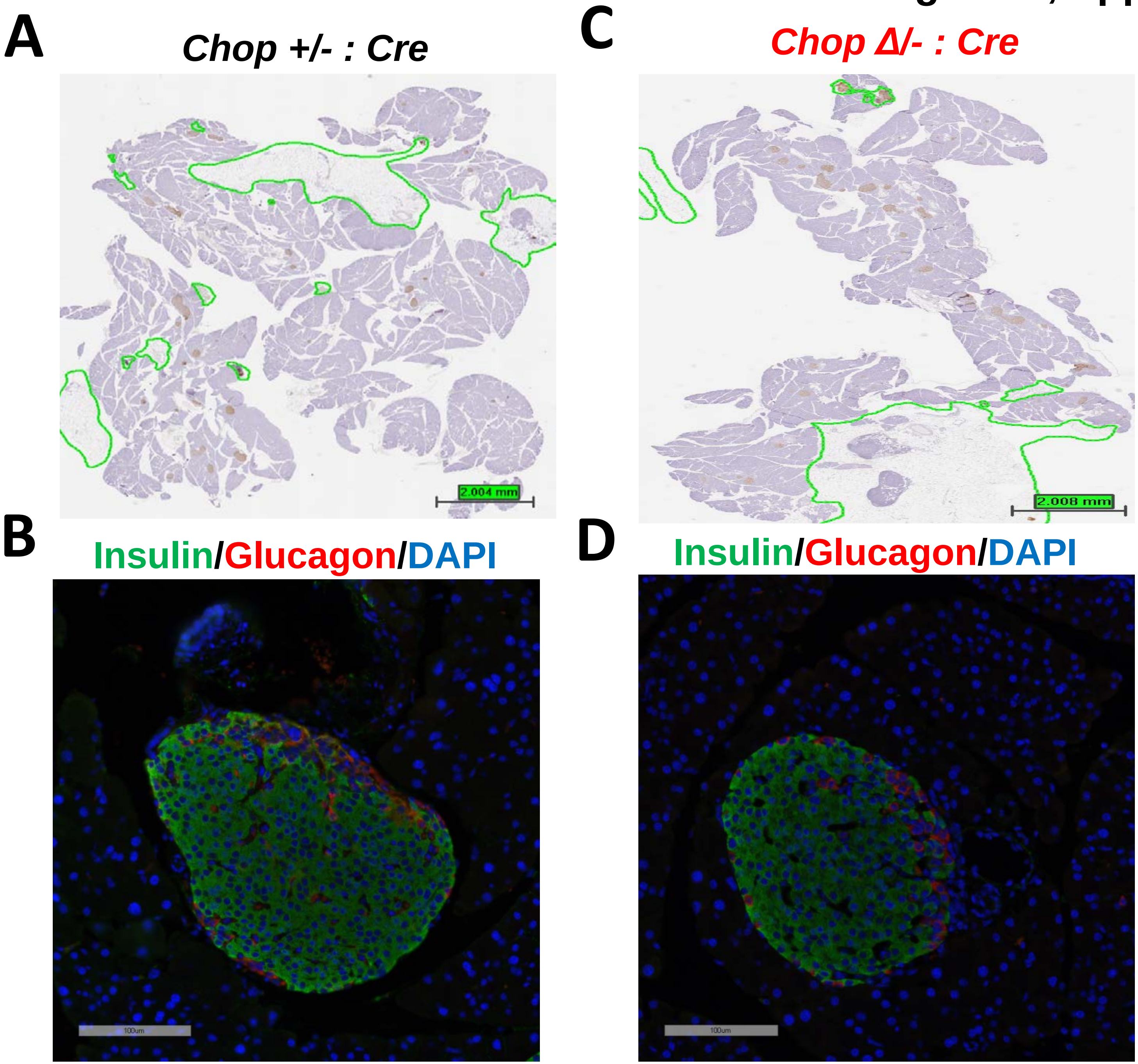

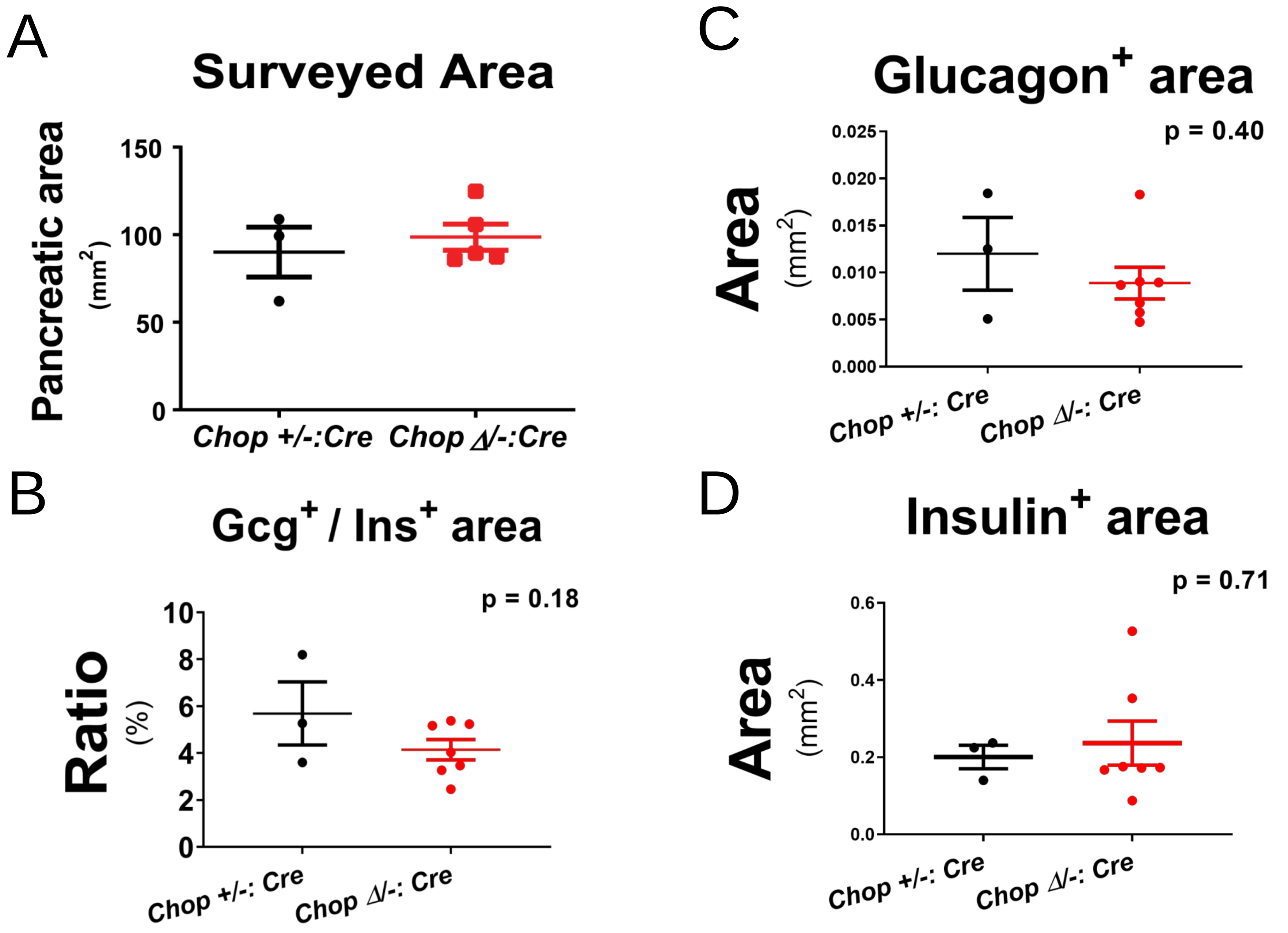
A
Surveyed Area
Pancreatic area
(mm2)
150
100
50
0
Chop +/-:Cre
Chop Δ/-:Cre
B
Gcg+ / Ins+ area
p = 0.18
Ratio
(%)
10
8
6
4
2
0
Chop +/-: Cre
Chop Δ/-: Cre
C
Glucagon+ area
p = 0.40
Area
(mm2)
0.025
0.020
0.015
0.010
0.005
0.000
Chop +/-: Cre
Chop Δ/-: Cre
D
Insulin+ area
p = 0.71
Area
(mm2)
0.6
0.4
0.2
0.0
Chop +/-: Cre
Chop Δ/-: Cre

Yong et al., Sppl. Figure S6

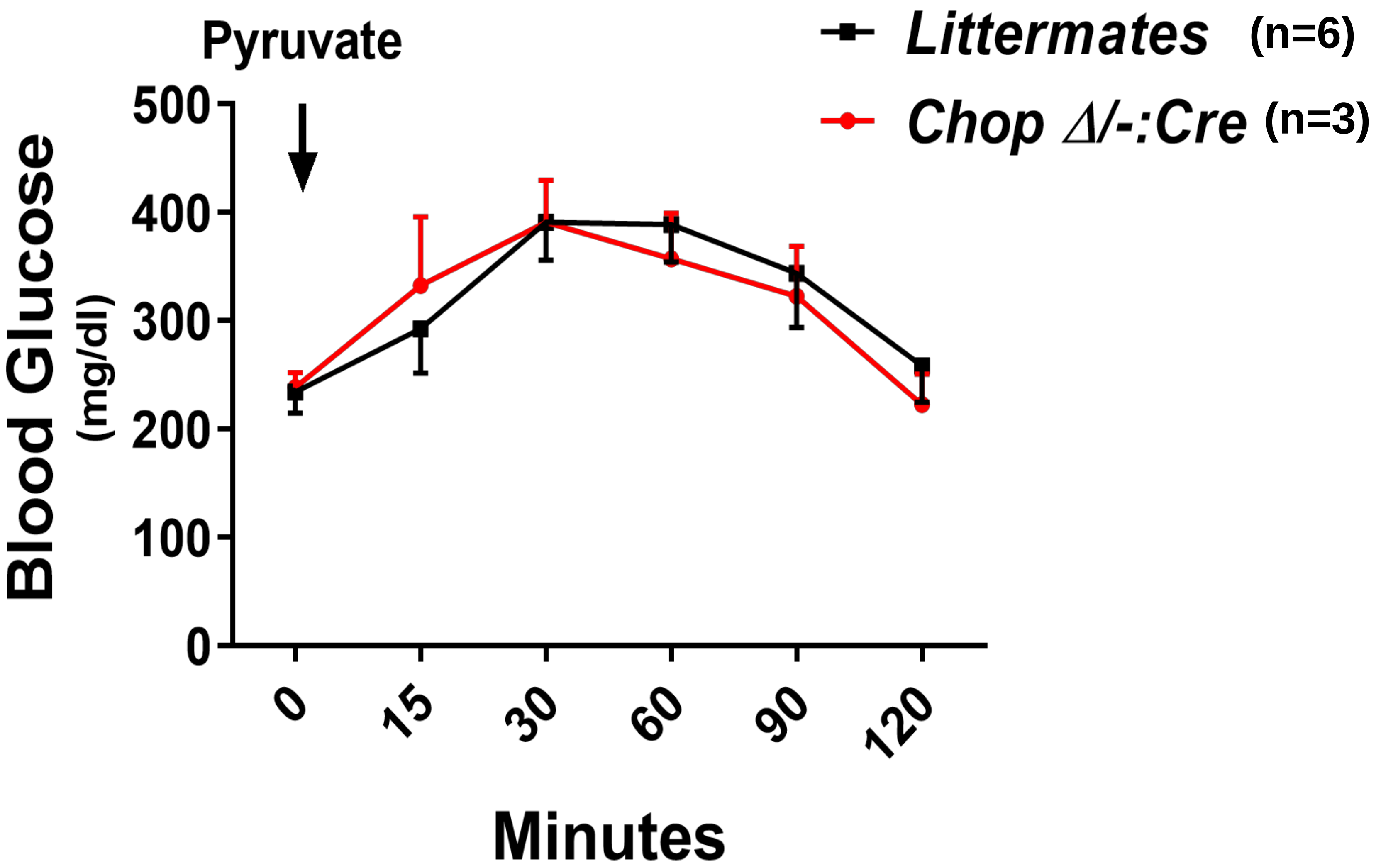

Yong et al., Sppl. Figure S7

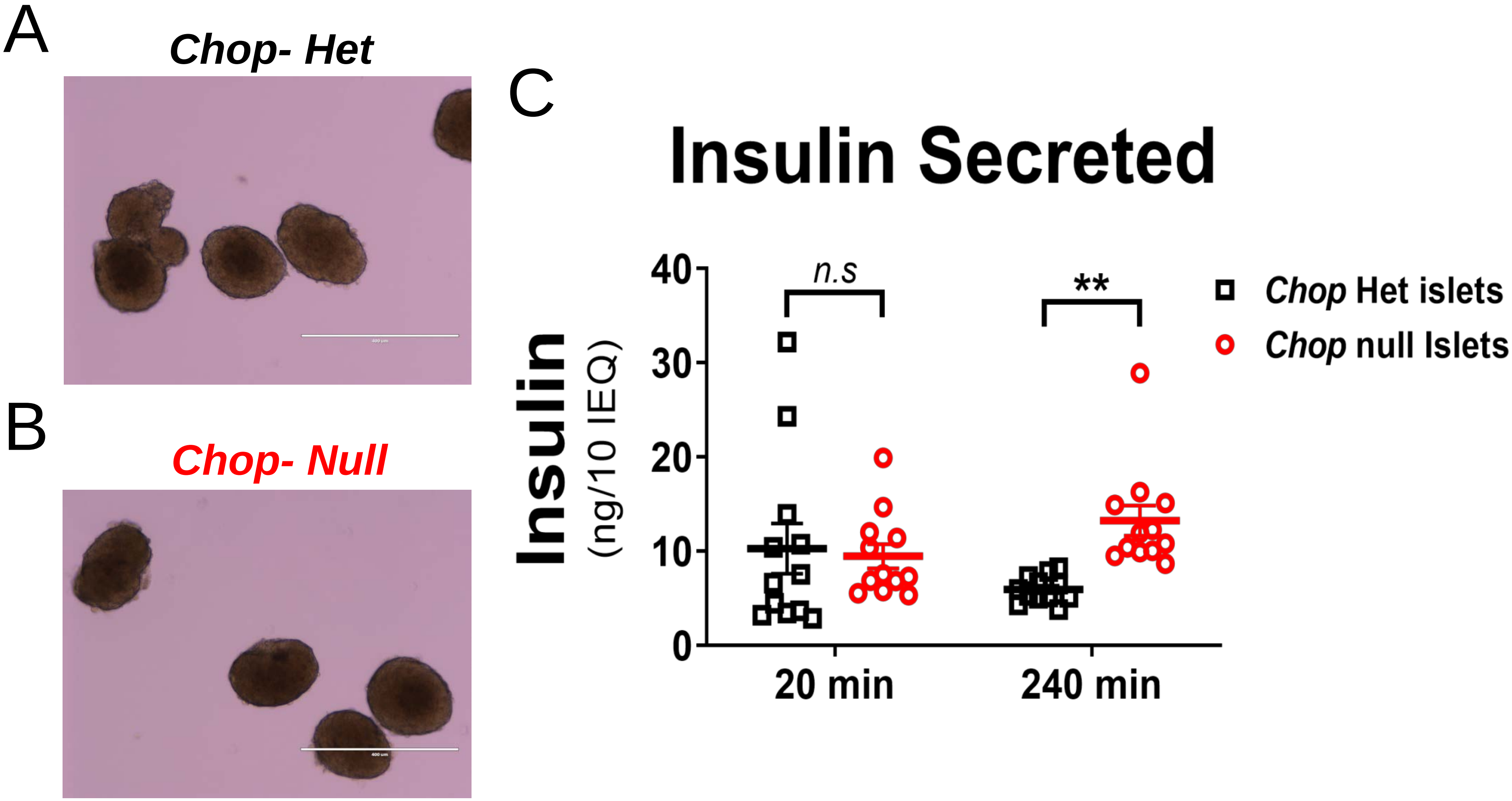

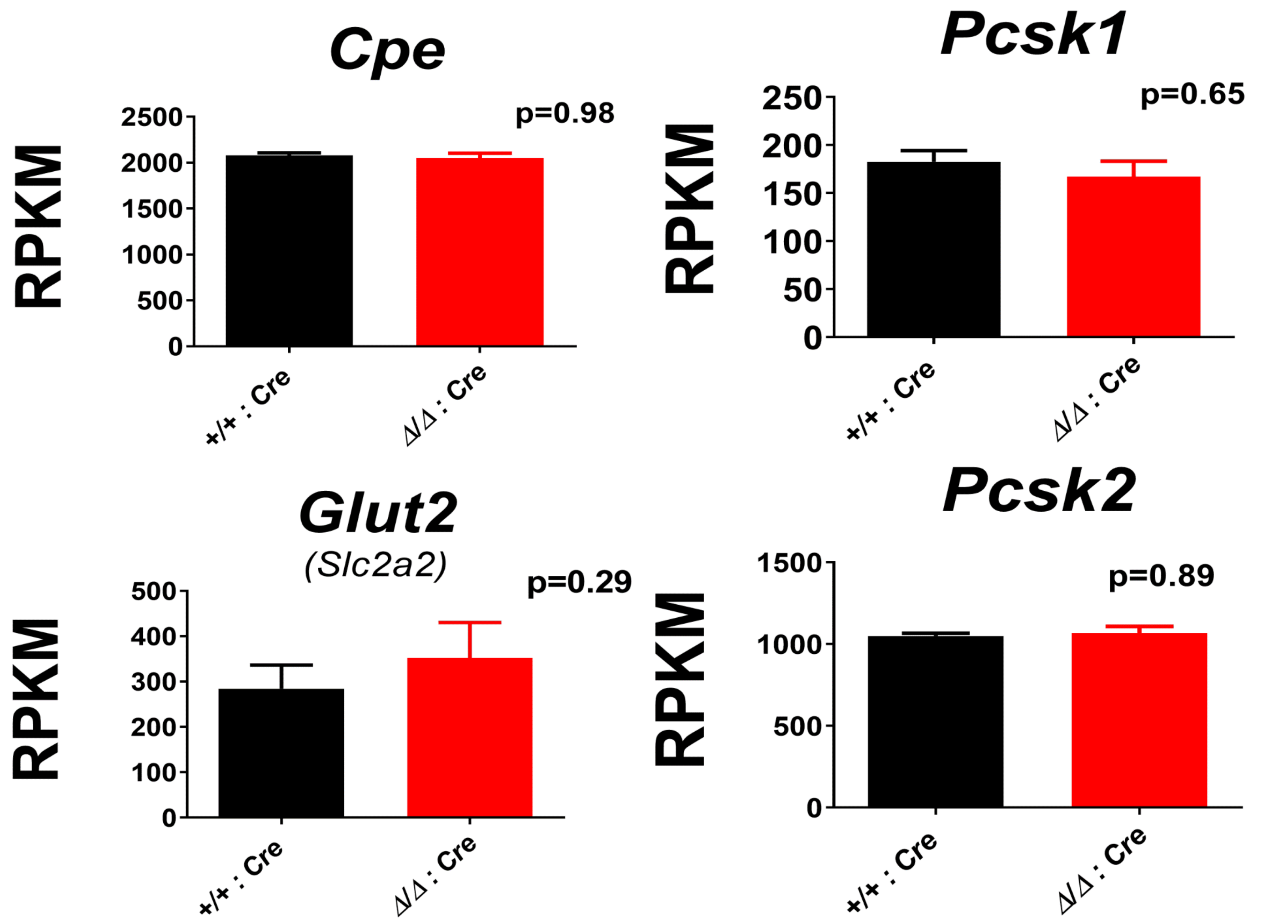
Cpe
RPKM
2500
2000
1500
1000
500
0
p=0.98
+/+ : Cre
Δ/Δ : Cre
Pcsk1
RPKM
250
200
150
100
50
0
p=0.65
+/+ : Cre
Δ/Δ : Cre
Glut2
(Slc2a2)
RPKM
500
400
300
200
100
0
p=0.29
+/+ : Cre
Δ/Δ : Cre
Pcsk2
RPKM
1500
1000
500
0
p=0.89
+/+ : Cre
Δ/Δ : Cre

Yong et al., Sppl. Figure S9

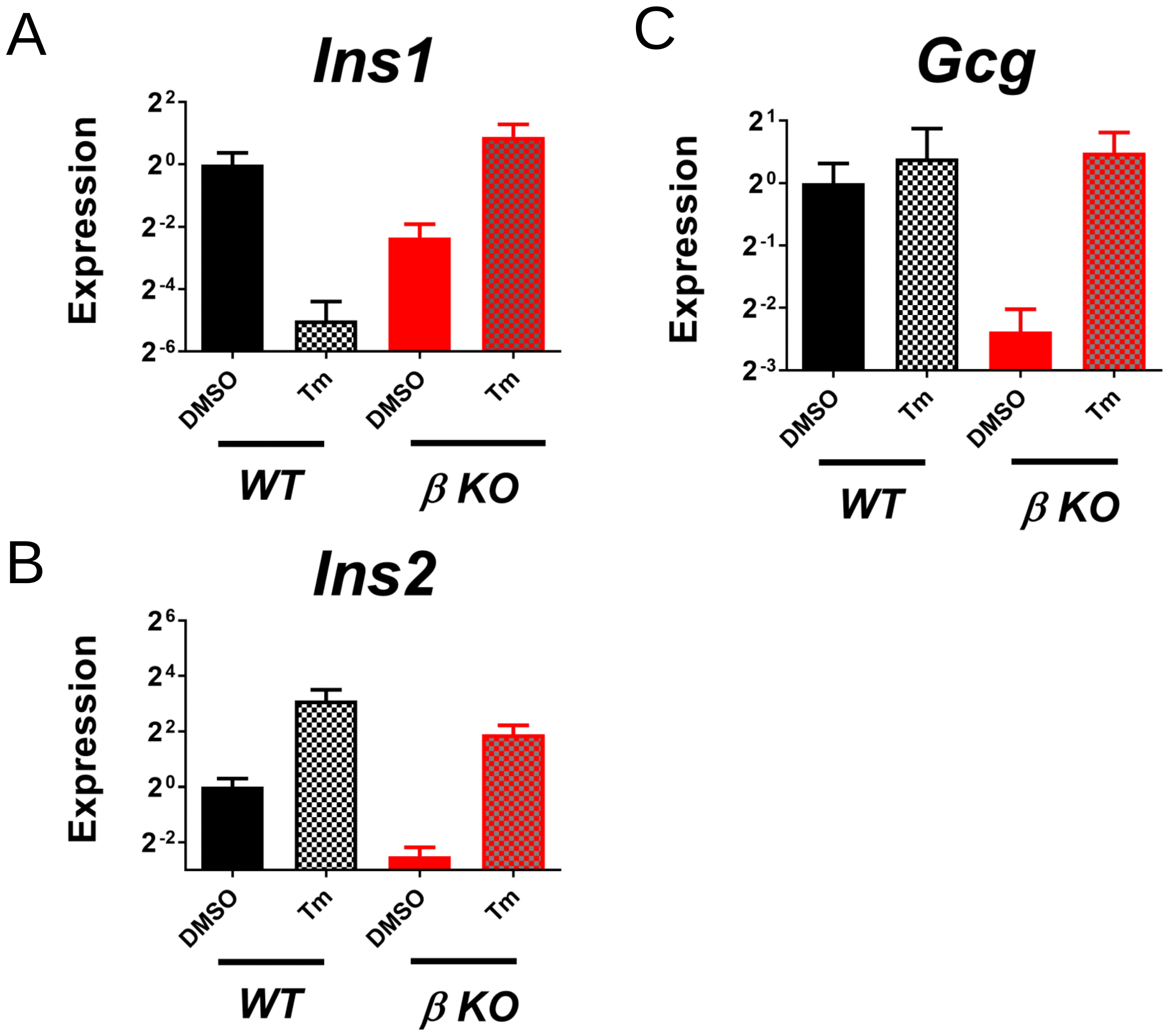

#### KEY RESOURCES TABLE

| REAGENT or RESOURCE | SOURCE | IDENTIFIER |
|---|---|---|
| **Antibodies** | | |
| Anti-Glucagon | Cell Signaling Tech. | 2760S |
| Anti-Insulin, for IP | Stuart Woodhead, personal gift | Invitron (3B1) |
| Anti-Insulin, for Western blot | Santa Cruz BioTech. | Sc-9168 |
| Anti-Insulin, for IHC and IF | Thermo Scientific | PA1-26938 |
| Anti-BiP | Linda Hendershot, personal gift | N.A. |
| Anti-Actin | MP Biomedicals | 691001 |
| **Bacterial and Virus Strains** | | |
| N.A. | | |
| **Biological Samples** | | |
| N.A. | | |
| **Chemicals, Peptides, and Recombinant Proteins** | | |
| Collagenase P | Roche | 11213873001 |
| Humulin | Eli Lilly | Humulin R U-100 |
| Histopaque -1119 | Sigma-Aldrich | 11191-100ML |
| Histopaque -1077 | Sigma-Aldrich | 10771-100ML |
| Halofuginone | Fisher Scientific | 5057630001 |
| EasyTag Express | PerkinElmer | NEG 772007MC |
| Tunicamycin | Santa Cruz Biotechnology | sc-3506 |
| Tamoxifen | Sigma-Aldrich | T5648 |
| Fura | Millipore-Sigma | 344911 |
| Cyclopiazonic acid | Sigma | C1530 |
| **Critical Commercial Assays** | | |
| Mouse Insulin ELISA kit | Mercodia | 10-1247-01 |
| Mouse C-peptide ELISA kit | Alpco | 80-CPTMS-E01 |
| Infinity Triglyceride kit | Thermo Scientific | TR22421 |
| Blood Glucose Test Strips | Embrace | APX02ABX0202 |
| RNeasy Kit | Qiagen | 74104 |
| **Experimental Models: Organisms/Strains** | | |
| Mouse, *Chop* gene deleted (germ-line) | The Jackson Laboratory | #005530, B6.129S(Cg)-Ddit3tm2.1Dron/J |
| Mouse, *Chop* gene floxed | The Jackson Laboratory | #030816, B6.Cg-Ddit3tm1.1Irt/J |
| Mouse, *RIP-CreER* transgenic | The Jackson Laboratory | # 008122, Tg(Ins2-cre/ERT)1Dam/J |
| **Rodent Diets** | | |
| Laboratory Mouse Diet | Teklad Global Irradiated Diet | # T.2918.15 |
| High Fat Diet (45% fat, kcal%) | Open Source Diets | # D12451 |
| **Oligonucleotides** (5'- to -3') | | |
| CTG CCT TTC ACC TTG GAG AC | IDT | *Chop-Ex3 Fwd* |
| CGT TTC CTG GGG ATG AGA TA | IDT | *Chop-Ex3 Rev* |
| CCA GCT ATA ATC AGA GAC CAT CA | IDT | *Ins1 Fwd* |
| GTT TGA CAA AAG CCT GGG TG | IDT | *Ins1 Rev* |

| | | |
|---|---|---|
| GGA GCG TGG CTT CTT CTA CA | IDT | *Ins2 Fwd* |
| GGT CTG AAG GTC ACC TGC TC | IDT | *Ins2 Rev* |
| GAT CTG GCA CCA CAC CTT CT | IDT | *b-Actin Fwd* |
| GGG GTG TTG AAG GTC TCA AA | IDT | *b-Actin Rev* |
| AAG AAC ACG CTT GGG AAT GG | IDT | *Xbp1 Fwd* |
| ACT CCC CTT GGC CTC CAC | IDT | *Xbp1 Rev* |
| GAG TCC GCA GCA GGT G | IDT | *sXbp1 Fwd* |
| GTG TCA GAG TCC ATG GGA | IDT | *sXbp1 Rev* |
| ATG GCC GGC TAT GGA TGA T | IDT | *ATF4 Fwd* |
| CGA AGT CAA ACT CTT TCA GAT CCA TT | IDT | *ATF4 Rev* |
| GGT GCA GCA GGA CAT CAA GTT | IDT | *BiP Fwd* |
| CCC ACC TCC AAT ATC AAC TTG A | IDT | *BiP Rev* |
| TGT CTA CAC CTG TTC GCA GC | IDT | *Gcg Fwd* |
| TGC CTT GCA CCA GCA TTA T | IDT | *Gcg Rev* |
| **Recombinant DNA** | | |
| N.A. | | |
| **Software and Algorithms** | | |
| Prism 7.0 | GraphPad Software | Version: 7 |
| Adobe Illustrator | Adobe | Version: CS 5.1 |
| Aperio ImageScope | Leica BioSystems | Version: 12 |
| CIMminer | NIH - NCI | https://discover.nci.nih.gov/cimminer |
| CFX Manager software | Bio-Rad | Version: 3.1 |
| Metafluor software | Molecular Devices | |
| Igor Pro software | WaveMetrics Inc | |
| **Deposited Data (at https://www.ncbi.nlm.nih.gov/sra by NCBI)** | | |
| RNA-Seq data for *Chop +/+: Cre* male mice, n=3 | SBP Analytical Genomics Core | *SRR10360208 /SRR10360209 /SRR10360210* |
| RNA-Seq data for *Chop Δ/Δ: Cre* male mice, n=3 | SBP Analytical Genomics Core | *SRR10360205 /SRR10360206 /SRR10360207* |
| RNA-Seq data for GLP1-ASO treated male mice, n= 3 - 4 | USCD Genomics core | SRR10350301 to SRR10350307 |
| **Other** | | |
| Fluorescent Image Analyzer | FUJIFILM | Model: FLA-5100 |
| CFX384 Touch™ Real-Time PCR Detection System | Bio-Rad | Model: 1855485 |

#### Materials and Methods

##### Generation and genotyping of *Chop*$^{Fe/Fe}$ (*Ddit3* $^{Fe/Fe}$) mice

The transgenic mice were generated in house, and characterized as previously published (*1*). This line is donated to The Jackson Laboratory, with strain catalogue #030816 ("*B6.Cg-Ddit3*$^{tm1.1Irt/J}$"). For genotyping of the mice carrying the floxed *Chop*/*Ddit3* allele, primers targeting the L83 element were used, as described on the website: [https://www.jax.org/strain/030816](https://www.jax.org/strain/030816).

##### Breeding strategy to generate the *Chop* β-KO mice

*Chop*$^{Fe/Fe}$ mice were first bred with a *RIP-CreER* transgenic mouse in which the fusion protein of Cre recombinase and a mutant human Estrogen Receptor (*ERT2*) is driven by the rat insulin II (Ins2) promoter (*2*) to generate the *Chop*$^{+/Fe}$: *RIP-Cre* founders. The *Chop*$^{+/Fe}$: *RIP-Cre* mice were further bred with *Chop*$^{+/Fe}$ mice to generate littermates of the following genotypes: 1). *Chop*$^{Fe/Fe}$: *RIP-Cre*; 2). *Chop*$^{+/Fe}$: *RIP-Cre*; 3). *Chop*$^{+/+}$: *RIP-Cre.* After TAM injections, specifically in the β-cells, the Ddit3 loci became: 1). *Chop*$^{\Delta/\Delta}$: *RIP-Cre*; 2). *Chop*$^{+/\Delta}$: *RIP-Cre*; 3). *Chop*$^{+/+}$: *RIP-Cre* respectively.

In selected experiments, *Chop*$^{-/-}$ mice were used (instead of *Chop*$^{+/Fe}$ mice) to breed with the *Chop*$^{+/Fe}$: *RIP-Cre* founder so that progeny of the following genotypes can be generated: 1). *Chop*$^{Fe/-}$ : *RIP-Cre*; 2). *Chop*$^{+/-}$ : *RIP-Cre*. After TAM injections, specifically in the β-cells, the Ddit3 loci became: 1). *Chop*$^{\Delta/-}$: *RIP-Cre*; 2). *Chop*$^{+/-}$: *RIP-Cre*; respectively.

##### Mouse diet and husbandry

Standard mouse diet (Teklad Global Irradiated Diet # T.2918.15) and water were available *ad libitum* at all times, except for a 4 to 6-h fasting period before intraperitoneal glucose tolerance test (IPGTT) when indicated. For the high fat diet studies (HFD, 45% fat in Kcal%, Open Source Diets # D12451), male mice from 2-3 litters were mixed and matched, with n= 2-4 mice per cage without genotype separation. Mice were only separated into individual cages for food intake measurement, when indicated. All procedures were performed by protocols and guidelines approved by the Institutional Animal Care and Use Committee (IACUC) at the SBP Medical Discovery Institute (AUF 17–066).

##### Tamoxifen injections

Deletion of the floxed *Chop* allele in pancreatic β-cells was induced by intraperitoneal injection of tamoxifen (TAM, Sigma Cat # T5648) dissolved in 10% ethanol in PBS in the dose of 2 mg per injection. Four to five doses of 4 - 10 mg TAM were administered over 2 weeks to adult mice (*2, 3*).

##### Homeostatic Model Assessment of β-cell function and IR calculation

The β cell function was calculated using the equation HOMA-β = [(360 x Insulin)/(Glucose -63)]%, and the IR was calculated using the equation HOMA-IR = (Glucose x Insulin/ 405), using the measurements of blood glucose concentration expressed in mg/dL and serum insulin concentration expressed in µU/mL, taken from mice after 4-6 hr fasting.

##### Targeted delivery of antisense oligonucleotides to pancreatic β-cells

For effects of Chop gene silencing in β-cells, *Chop* gene floxed mice (*B6.Cg-Ddit3*$^{tm1.1Irt/J}$, Jackson Laboratory Cat. #030816) were treated twice with GLP1-Chop ASO, once every 5 to 7 days, by subcutaneous injections of PBS diluted GLP1-ASO compounds in a volume of 0.2 mL per injection. GLP1-control ASO was administered to control littermate mice in parallel at the same time. A method describing the synthesis and characterization of the GLP1-ASO moiety can be found in Ammala *et al.* (*4*). Three days after the second GLP1-ASO injection, mice were sacrifice for islets isolation and characterization.

##### Nascent protein labeling and ProIns/Ins abundance analysis

Primary islets were isolated and incubated overnight in RPMI medium supplemented with 10% fetal bovine serum and 1% Pen/Strep antibiotics mix, before usage. A total of 40 medium-sized islets were used for each sample in the experiment. After pre-incubation in a tissue culture incubator for 15 min in the labeling medium (containing no methionine/cysteine, Gibco Cat # 21013-024), medium was replaced with 0.3 mL of labelling medium supplemented with 150 µCi/mL of [$^{35}$S]-cysteine/methionine (EasyTag Express, PerkinElmer Cat # NEG 772007MC) for 20 min, for pulse labelling of newly synthesized ProIns and Ins. Duplicate samples of both genotypes were harvested at the end of pulse labeling to detect labeling efficiency, while other samples were further incubated for a chase period of 1 hr or 2 hrs, respectively. Medium were replaced with complete DMEM medium containing 300 mg/dL of glucose with un-labeled cysteine/methionine during the chase period. Islets were harvested after two washes with cold DMEM medium before they were lysed in cell lysis buffer containing 50 mM Tris (pH 7.4), 150 mM NaCl, supplemented with 1% Triton X-100 (v/v) and 0.1% deoxycholate (v/v). Islets lysates were separated on a SDS-PAGE gel (12% Criterion XT Bis-Tris Protein Gel; Bio-Rad # 345-0118), and radiolabeled ProINS+INS bands were visualized on the dried gel using a Phosphor Screen on a Fluorescent Image Analyzer (FUJIFILM, Model FLA-5100).

To detect insulin secreted into the medium, conditioned DMEM media were collected at the same time when radiolabeled islets were harvested, and were frozen at -80 ºC until the day of assay. Conditioned media were diluted 80 fold with fresh DMEM media, and 10 µL of the diluted sample was used on an anti-Mouse Insulin ELISA plate (Mercodia, #10-1247-01) for insulin concentration determination.

##### Insulin Sensitivity Test

Mice were fasted for 4-5 hrs, followed by an i.p. injection of insulin (0.75 U/kg body weight of Humulin, Eli Lilly). Blood samples were collected through the tail vein, with the blood glucose levels measured using a *OneTouch Ultra* glucometer (LifeScan, Inc).

##### Liver TG concentration determination

TG concentrations in fresh liver homogenates were determined by a commercial TG kit (Thermo Scientific, Cat # TR22421) after tissue homogenization and sodium deoxycholate treatment (0.5% end concentration, weight/volume). TG content were further standardized by the wet weight of the liver tissue.

**Quantitative RT-PCR (qRT-PCR) assays**

RNA was extracted from islets using RNeasy (Qiagen, Cat # 74104). The relative amounts of mRNAs were calculated from the comparative threshold cycle (Cq) values relative to β-actin. Nucleotide sequences for used primers are shown in the *Key Resources Table*. Data was analyzed using the normalized Gene Expression (*ΔΔ*Cq) method offered by Bio-Rad CFX Manager software (Version: 3.1).

**Hyperglycemic Clamp Assay**

Hyperglycemic clamps were conducted in conscious mice as previously described (*5*). Briefly, mice were implanted with catheters into the right jugular vein and left carotid artery for infusions and sampling, respectively. Following a 5-day recovery, hyperglycemic clamps were conducted in 5h-fasted mice. 50% dextrose was infused at a variable rate to maintain glucose levels at ~300 mg/dL. Blood glucose (5 μl) was measured at $t = -15, -5, 5, 10, 15$, and 20 min and then every 10 min until $t = 120$ min. Samples (50 μl) to measure plasma insulin and C-peptide were taken at $t = -5$, 5, 10, 15, 20, 40, 80 and 120 min.

**$Ca^{2+}$ imaging assay on primary islets**

Free $Ca^{2+}$ was measured as previously described (*6*). Excitation was provided using an LED-based excitation source (pE-340fura; CoolLED, UK). The excitation (x) or emission (m) filters (Chroma Technology, Bellows Falls, VT, USA) used were as follows: Fura-2, 340/10x and 380/10x , 535/30m (R340x/380x; 535m). Fluorescence emission was collected using an ORCA-Flash4.0OLT camera (Hamamatsu, Japan) at 6s intervals. Data were acquired and analyzed using *MetaFluor* software (Molecular Devices, Sunnyvale, CA, USA), and plotted using *Igor Pro* software (WaveMetrics Inc., Lake Oswego, OR, USA). All statistical analysis was done using *Prism* software (GraphPad, La Jolla, CA, USA).

**Whole-genome analysis of messenger RNA expression profile in islets**

Complementary DNAs were prepared for mRNA-seq according to the manufacturer's instructions (*Illumina*). cDNA fragments were prepared for Next-Generation Sequencing as described previously (*7*). Islets from three male mice were prepared for both genotypes, with the number (proportion) of successfully aligned reads from a sample ranging from 55 million to 80 million (> 98% mapping rate). *TopHat Splice-aware Aligner* was employed to align reads to the mouse reference genome (*mm10*) plus known splice junctions, created by ERANGE scripts and UCSC known genes (http://genome.ucsc.edu). Counts of reads were normalized to RPKM (reads per kilobase pair per million reads mapped) values for transcript abundance comparison. Trimmed Mean of M (TMM) values were used for between-sample comparisons. For differential expression test, Generalized Linear Model likelihood ratio test was implemented in EdgeR software. For GLP1-ASO treated samples, RNA extracted from mouse islets using RNeasy (Qiagen, Cat # 74104) was further prepared using Illumina's TruSeq library prep kit. RNA-Seq data analysis was performed by *Sanford Burnham Prebys Bioinformatics Core* in a similar workflow. Adapter remnants of sequencing reads were removed using *Cutadapt* (version 1.18) (*8*). RNA-Seq sequencing reads were aligned using *STAR aligner* (version 2.7) (*9*). Mouse genome version 38 and Ensembl gene annotation version 84 were used in the alignment and quantification. Transcripts per Million (TPM) and read counts were quantified using RSEM (version 1.3.1) (*10*). Estimated read counts from RSEM were compared using the R Bioconductor package *DESeq2* following generalized linear model based on negative binomial distribution (*11*). Genes with Benjamini-Hochberg–corrected p value< 0.05 and fold change >=2 or <=0.5 were selected as significantly differentially expressed genes.

**Heat-map generation.**

Color-coded Clustered Image Maps (CIMs) (or "heat maps") to visualize expression levels for individual genes of interest. A heat map was generated using the CIMminer (https://discover.nci.nih.gov/cimminer). *Minkowski* distance was applied to cluster genes of similar changes in the expression level.

### Table I. Breeding Strategy – Examples

| Genotypes For Breeding Pair | Control Littermate Genotype (Expected Ratio for Male) | *Chop-βKO* Genotype (Expected Ratio for Male) | Used in Figures |
|---|---|---|---|
| ***Chop Fe/Fe : RIP-Cre*** **X** ***Chop Fe/Fe : 0*** | ***Chop Fe/Fe : RIP-Cre*** ***+ Diluent injections*** (½ x ½ = 1/4) | ***Chop Fe/Fe : RIP-Cre*** ***+ TAM injections*** (½ x ½ = 1/4) | **Figure 1** (1A to 1D) |
| ***Chop Fe/+ : RIP-Cre*** **X** ***Chop Fe/+ : 0*** | ***Chop +/+ : RIP-Cre*** (½ x ½ x ¼ = 1/16) | ***Chop Fe/Fe : RIP-Cre*** (½ x ½ x ¼ = 1/16) | **Figure 1** (1F to 1L) |
| ***Chop Fe/Fe : RIP-Cre*** **X** ***Chop +/- : 0*** | ***Chop Fe/+ : RIP-Cre*** (½ x ½ x ½ = 1/8) | ***Chop Fe/- : RIP-Cre*** (½ x ½ x ½ = 1/8) | **Figure 2** (2A to 2D) |
| ***Chop -/- : RIP-Cre*** ***X*** ***Chop Fe/+ : 0*** | ***Chop +/- : RIP-Cre*** (½ x ½ x ½ = 1/8) | ***Chop Fe/- : RIP-Cre*** (½ x ½ x ½ = 1/8) | **Figure 2** (2F to 2K) |

#### Table II. Mice for Hyperglycemic Clamp Study

| **Genotype** | **Male** | **Female** | **Total** | Body Weight (g) | Fasting blood glucose (mg/dL) |
|---|---|---|---|---|---|
| *Chop +/- : RIP-CreER* | 2 | 1 | 3 | 36.8 ± 1.2 | 173 ± 12 |
| *Chop Δ/- : RIP-CreER* | 6 | 2 | 8 | 39.6 ± 2.9 n.s, # | 197 ± 26 n.s, ## |

Note:

Data are displayed as Mean ± S.E.M.

#: $p$ = 0.58 by two-tailed t-test, compared to the control group.

##: $p$ = 0.59 by two-tailed t-test, compared to the control group.
